# Xeno amino acid alphabets form peptides with familiar secondary structure

**DOI:** 10.64898/2026.01.28.701962

**Authors:** Sean M. Brown, Tadeáš Kalvoda, Robin Kryštůfek, Ján Michael Kormaník, Mikhail Makarov, Václav Verner, Rozálie Hexnerová, Pavel Srb, Erik Andris, Lucie Bednárová, Markéta Pazderková, Martin Lepšík, Jan Řezáč, Jan Konvalinka, Václav Veverka, Lubomír Rulíšek, Stephen Freeland, Klára Hlouchová

## Abstract

The central dogma of molecular biology describes how genetic information stored in nucleic acids guides the formation of structured proteins from a single molecular alphabet of twenty amino acids (C20)^1,2^. The extent to which C20 is uniquely capable of forming structural polymers remains a fundamental open question. Here we demonstrate that peptides built from other, “xeno” amino acid alphabets can adopt protein-like secondary structural motifs. Having designed two different xeno alphabets, with them we constructed both combinatorial random sequence libraries and specific, designed 25-mer sequences. We report sequence-dependent structural motifs that result according to circular dichroism, infrared spectroscopy and nuclear magnetic resonance interpreted by molecular dynamics simulations, supported by extensive quantum mechanical calculations. This evidence, including a solution-phase NMR-resolved helical motif, demonstrates that the potential for amino acids to form structure bearing sequences is not unique to life’s alphabet, and thereby reveals a previously unexplored sequence-structure space. Results inform the search for extraterrestrial life, lay foundations for incorporating novel synthetic functional groups and heteroatoms into structure-bearing alphabets, and provide the first truly independent data with which to test and improve all that has been learned about protein folding from the study of life’s 20 side chains.

## Introduction

For more than 3.5 billion years, all life has shared a common molecular architecture, formally referred to as the Central Dogma of molecular biology^1^: cells translate genetic information, encoded by a universal set of nucleotides, into proteins built from a set of twenty canonical amino acids (C20). As a sequence of amino acids link together through peptide bonds, it is the order of differing sidechains which guides the three-dimensional structure, and therefore function, of the resulting protein^2^. Such deep, shared features of life raise important questions: does life’s remarkable persistence, its successful diversification to indefinite environmental limits^3^, perhaps even its very existence^4^, result from this specific collection of nucleotides and amino acids, or are the chemical details which define biology just one of many possible alternatives?

Evidence increasingly favours the latter interpretation by showing that one half of life’s central dogma, namely the nucleic acids, can be reconfigured into other viable options. Several alternatives have been designed and used to produce structured and catalytic variants of life’s genetic polymers^5–9^. Exploration has gone beyond changes to the genetic letters (nucleobases) by testing alternatives to the sugar phosphate backbone^10^. On the other side of the genetic code, recent experimental evidence now validates a large body of theoretical work suggesting that the canonical amino acid alphabet likely evolved from a simpler, smaller set (Gly, Ala, Asp, Val, Ser, Glu, Pro, Leu, Thr, Ile)^11–13^. This foundational set of 10 amino acids, commonly known as the ‘Early 10’ (E10), can form compact, soluble structures without the aid of chaperone proteins that evolved subsequently^14–18^. Structure-bearing peptides built with E10, a subset of C20, currently represents the largest deviation from C20 known to science. This empirical data for biochemical flexibility is consistent with theory to suggest that natural selection honed the canonical genetic molecules as a local or even global optimum within a much larger chemical space of possibilities^10,19–21^.

The mere existence of alternative molecular architectures broadens scientific understanding of how life could be engineered in the laboratory and the aligned question of what we might expect from an independent origin of life beyond Earth^19^. For amino acids, hundreds of individual “xeno” amino acids not found within life’s genetic code (xAAs) have been experimentally incorporated into otherwise canonical proteins, often for biotechnological applications^22–25^. Most recently, a pioneering search has discovered energetically favorable conformations, unknown from natural proteins, for dipeptide repeats that mix canonical and xeno amino acids^26^. At present however it is unknown whether any xeno set of amino acids could build structured proteins.

Here, we expand this exciting frontier by testing whether peptides bearing familiar secondary structure can be constructed from a very different set of amino acids than that which has enabled life as we know it. Given the theoretically infinite number of possible amino acid sets, the challenge is determining which amino acids to select. If the canonical alphabet (or its subset E10) represents a set of molecules shaped by natural selection, then its unusual physicochemical features could be abstracted into a guide for forming a novel xeno amino acid ‘alphabet’^27–29^. This study therefore combines recent progress incorporating individual xeno amino acids into canonical proteins with amino acid alphabet design theory and empirical evidence that smaller alphabets are possible to evaluate the structure-forming potential of alternative (xeno) alphabets similar to E10. Specifically we design two sets, each comprising 10 chemical structures unknown to biology. This was done by searching and analysing amino acid chemistry space^30^ for alternative sets which emulate the distribution of certain physicochemical properties of E10 (see Fig. 1). We then use density functional theory (DFT) methods together with implicit solvation models to carry out extensive conformational sampling of the individual amino acids within the newly designed sets, mapping their potential energy surfaces (PES) *ab initio*. The resulting candidate amino acid alphabets are filtered to two final sets based on a comparison with E10 in terms of PES. The selected alphabets (denoted Xeno 1 and Xeno 2) bear no compositional overlap with the canonical set, other than the simplest α-amino acid possible, glycine. Glycine can be viewed as a point of origin, an “identity element” or “zero-vector” in amino acid chemistry space. Secondary structure propensity is then measured for these xeno alphabets, as characterised within combinatorial peptide libraries, as well as for a panel of individual peptides. Results demonstrate the first experimentally verified structure-forming set of amino acids different from the one alphabet genetically encoded by all life on Earth.

**Figure 1.**
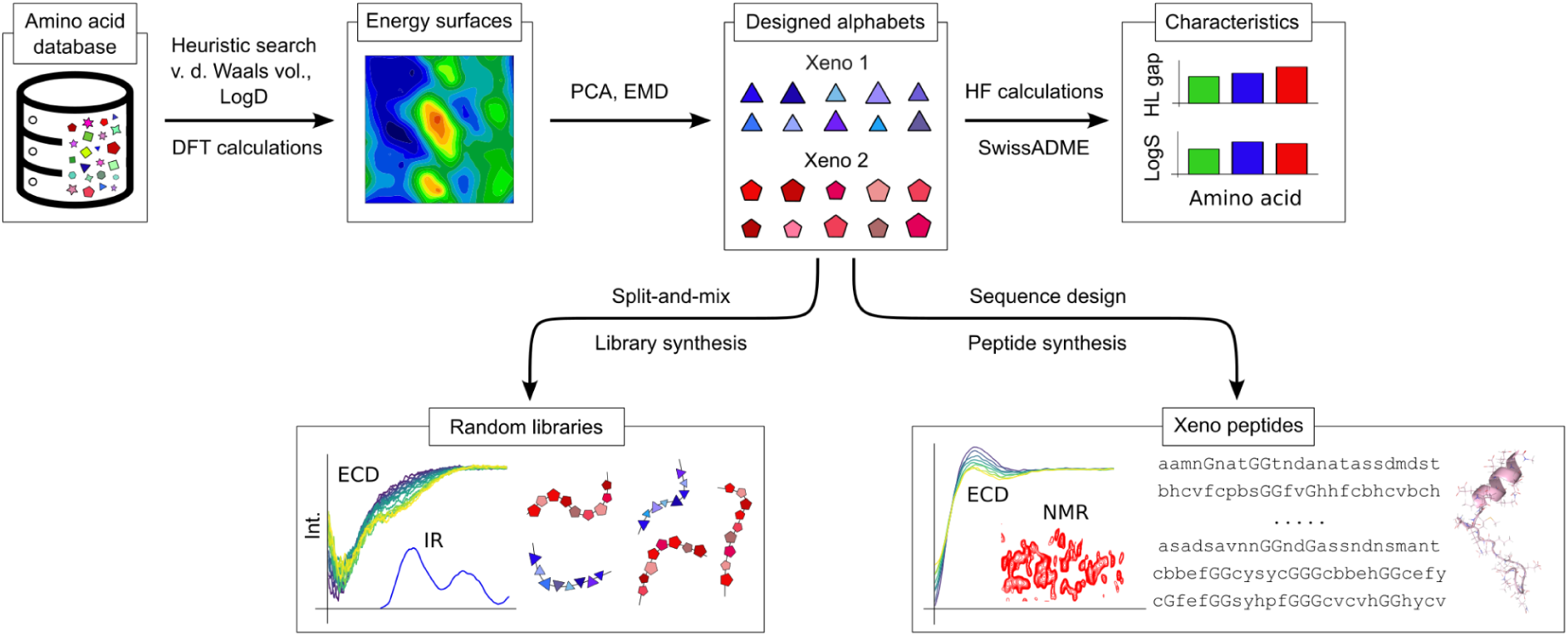
The experimental pipeline, from design to characterization, of peptides comprising the Xeno 1 and Xeno 2 amino acid alphabets. Design begins with a library of commercially available amino acids. The physicochemical properties (Van der Waals volume and hydrophobicity at pH 7) are then estimated for a xeno alphabet heuristic search. The potential energy surfaces for each amino acid which returns in the search is calculated and two xeno amino acid alphabets, Xeno 1 and Xeno 2, are selected based on post-search metrics (e.g. PCA and EMD). The similarity of these alphabets to the E10 is further assessed via calculation of HOMO-LUMO gaps and LogS: their ability to form secondary structures is measured *in vitro* with ECD, IR, and NMR spectroscopy for random combinatorial 25-mer libraries and for designed sequences.

## Results and Discussion

### Computational evaluation of the rationally-designed alphabets

Our rational design protocol (Extended Data Fig. 1) ultimately yielded two candidate xAA alphabets, Xeno 1 and Xeno 2 (Fig. 2a). Each alphabet was designed to emulate the physicochemical profile of E10 in terms of size, hydrophobicity, PES, and side chain protonation states. Both resulting xeno alphabets contain some amino acids that are polar in water at neutral pH, some that are non-polar, and some with unclear polarity at these conditions. Both Xeno 1 and Xeno 2 contain some amino acids structurally similar to the genetically encoded amino acids, such as homoglutamine, *O*-methyl-homoserine, or 3-(pyrazol-1-yl)-alanine. Indeed, both xeno alphabets contain some amino acids that are found within the collective metabolism of life and/or were widely available to life’s origins through prebiotic chemistry. Examples include 2-aminobutyric acid, γ-carboxyglutamic acid, ethionine, norleucine, norvaline, and selenomethionine^11,12,27,31–36^.

**Figure 2.**
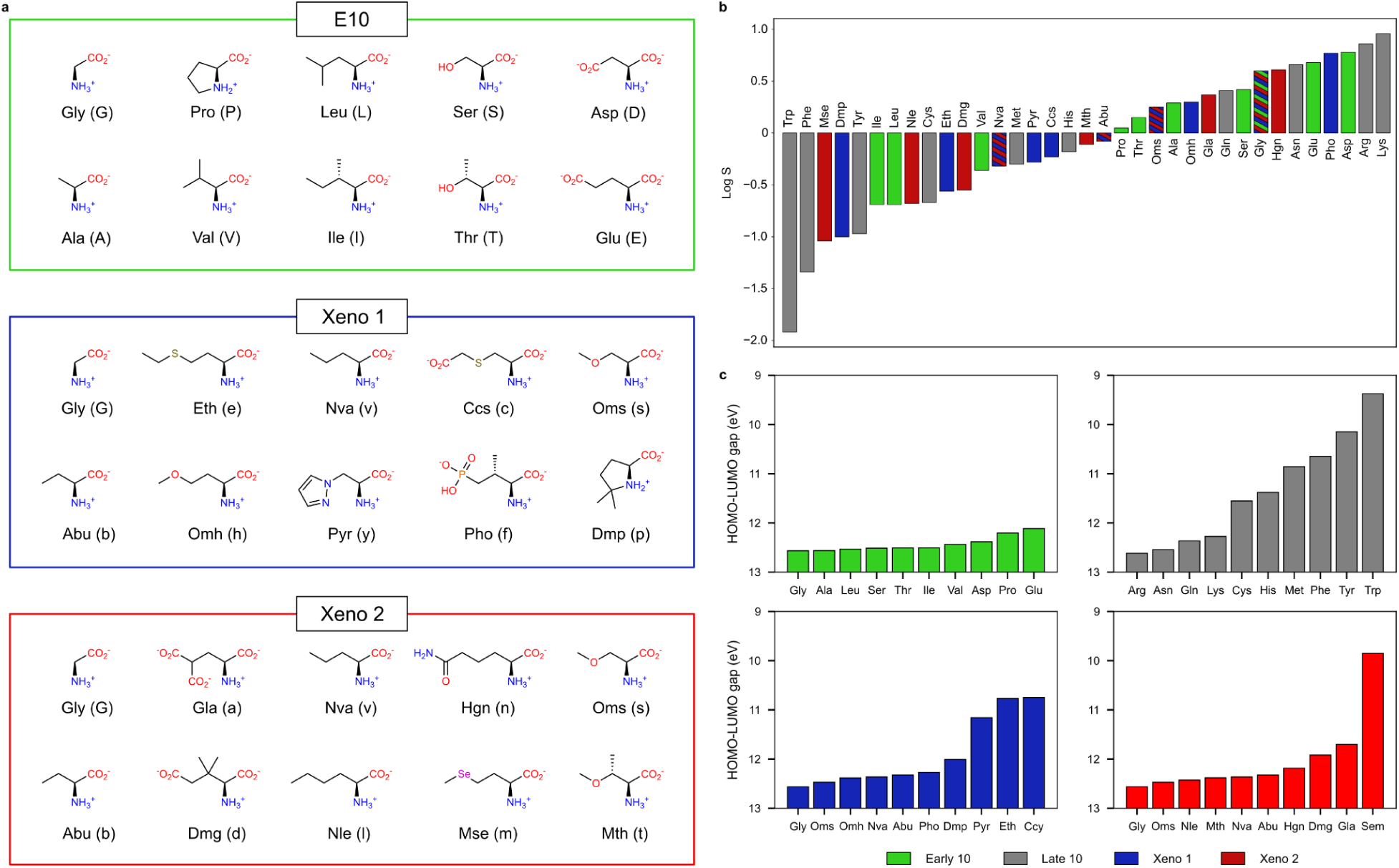
Composition and properties of designed xeno amino acid alphabets. **a,** The reference E10 alphabet and two selected xeno amino acid alphabets. Where standard one- and three-letter abbreviations exist, they are used; new abbreviations are proposed for all other amino acids. All amino acids are depicted in the preferred protonation state in water at pH 7. **b**, Estimation of log(S) of amino acids using the ESOL model for (i) E10 - green, (ii) the late 10 canonical amino acids - grey, (iii) Xeno 1 - blue, (iv) Xeno 2 - red. **c**, HOMO-LUMO gap energies, calculated at HF/def2-TZVPD/COSMO (*ε_r_*=80) for (i) the E10 alphabet - green, (ii) the late 10 canonical amino acids - grey, (iii) Xeno 1 - blue, (iv) Xeno 2 - red.

For quantitative comparison with E10 and C20, we estimated descriptors of solubility (logS; see methods), a key determinant of protein stability^37^, and HOMO–LUMO gaps, a widely applicable theoretical predictor of chemical reactivity^38–40^. In terms of logS, both Xeno 1 and Xeno 2 resemble the span and distribution of E10 (Fig. 2b): polar Hgn and charged Pho are almost as soluble as Glu and Asp, respectively, while non-polar Mse and Dmp are even less soluble than Ile and Leu. This similar solubility distribution likely reflects the rational design strategy to find alphabets with a similar LogD profile to that of E10 at neutral pH. Despite the fact all rational design was to emulate E10, the distributions of Xeno 1 and Xeno 2’s HOMO-LUMO gaps resemble C20. Understandably, a preponderance of amino acids in each alphabet have similar HOMO-LUMO gaps to E10 (Fig. 2c). The somewhat unexpected resemblance to C20 comes from the presence of chemical features not found within E10: Ccs and Eth, likely due to sulphur (both comparable to methionine in this respect), Mse due to selenium, and Pyr due to an aromatic ring. Together, these results indicate that the rational design of both xeno alphabets appears to be a success in that both Xeno 1 and Xeno 2 appear to emulate features of E10.

Shifting focus from comparison with E10 to comparison with each other, both xeno alphabets share three xAAs: Abu, Oms, and Nva. In terms of atomic composition and functional groups, Xeno 1 is more exotic (chemically diverse) than Xeno 2. The former includes two amino acids containing sulphur, one containing phosphorus, one with an aromatic heterocycle, and one proline derivative. In contrast, the only novel functional groups of Xeno 2 that extend beyond both E10 and C20 are a geminal dicarboxylic amino acid, an ether, and a sidechain containing selenium.

### Secondary structure analysis and characteristics of random combinatorial sequence libraries

To explore the potential of both xeno alphabets and E10 to form any type of secondary structure within random 25-mer libraries, we measured both room-temperature and temperature-dependent circular dichroism (CD) spectra, and infrared (IR) spectra (Fig. 3). In addition to secondary structure analysis, for each library, we investigated across a pH range the changes in solubility with UV-Vis absorption and fluorescence spectroscopy (Extended Data Fig. 2), and aggregation propensity with size-exclusion chromatography (Extended Data Fig. 3).

**Figure 3.**
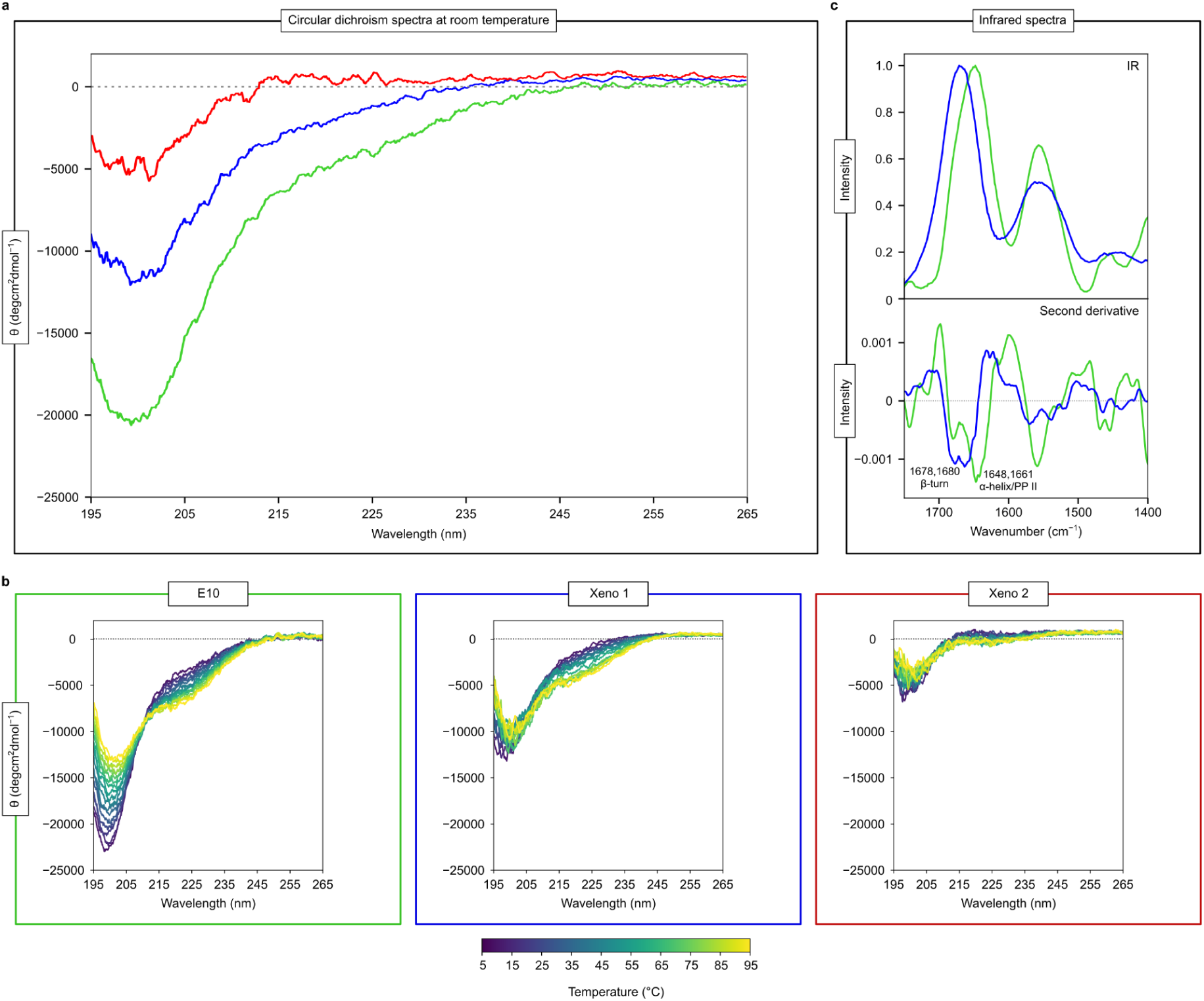
Experimental verification of xeno combinatorial libraries via CD and IR spectroscopy. **a,** Averaged circular dichroism spectra of 25-mer libraries at room temperature (20 °C) for E10 (green), Xeno 1 (blue), and Xeno 2 (red) **b,** Temperature dependence of CD spectra for E10 (green), Xeno 1 (blue), and Xeno 2 (red) combinatorial libraries. **c,** Infrared spectroscopy (IR) spectra and its second derivative for combinatorial Xeno 1 (blue) and E10 (green) libraries.

In the CD spectra observed at room temperature (Fig. 3a), all three peptide libraries exhibit a dominant negative band at ∼195 nm, indicating a predominance of disordered structural content^41–45^. Beyond this baseline of universal similarity, differences between the libraries’ CD spectra begin with the intensity of this spectral band, which is most pronounced for E10, intermediate for Xeno 1, and least intense for Xeno 2. In addition, both E10 and Xeno 1 display a negative shoulder at ∼220 nm. Together with the negative band at 195 nm, these spectral features are ambiguous to assign given that β-structures^42^, α-helices^42^, and disordered content^45^ manifest around this wavelength. In contrast, we observe a moderate positive signal around 215 nm in the Xeno 2 spectrum, suggesting a significant contribution from PPII helices^44–46^. The overall lower CD signal intensity for Xeno 1 and Xeno 2 compared to E10 could also indicate β-sheet content^41–44^. We therefore used temperature dependence CD and IR spectroscopy to gain further insight into the structural differences observed for the E10, Xeno 1, and Xeno 2 libraries.

Because temperature dependent CD monitors potential unfolding, it can differentiate more clearly between disordered or structured content in the libraries^44,45^. With increasing temperature, we observed a similar trend in the CD spectra of all libraries (Xeno 1, Xeno 2 and E10): the intensity of the spectral band at ∼198 nm decreased, accompanied by an increase in negative intensity at ∼220 nm with an isosbestic point at ∼210 nm (Fig. 3b). These spectral changes are characteristic of structural transition from PPII helix to disordered structure^44–46^. Temperature dependence CD thus supports the conclusion that both xeno libraries bear structural propensity that is most likely PPII helical structure. Xeno 1 spectral features resemble more closely those of E10 while the Xeno 2 library spectrum is likely affected by other additional structural elements (such as β-structures). The spectra of both Xeno libraries generally look very similar to those of short peptides comprising canonical (both E10 and C20) amino acids in both standard and temperature-dependent CD^15,41–43,46^.

To evaluate the potential presence of β-structures and possible β-aggregation, both of which are difficult to identify in CD spectra^43^, IR spectroscopy was employed for both E10 and Xeno 1 (the xeno library with higher resemblance to E10 in CD spectroscopy)^43,47–49^. In the IR spectra of E10 and Xeno 1 (Fig. 3c), no spectral bands characteristic of β-aggregation (∼1620 cm⁻¹) or of β-sheet structures (approximately 1623-1641 cm⁻¹ and 1674-1695 cm⁻¹) were detected. The spectral band observed at ∼1648 cm⁻¹ for E10 corresponds to either PPII, disordered, or α-helical structure^47–49^: IR alone cannot distinguish between these three interpretations and structure assignment is therefore possible here only when considering IR spectroscopy in combination with CD spectroscopy. The spectral band at ∼1680 cm⁻¹ in the IR spectra of both E10 and Xeno 1 may be attributed to β-turn structures^47–49^. The assignment of the spectral band at ∼1661 cm⁻¹ of Xeno 1 is more ambiguous to interpret as it lies at the extreme end of the range for both β-turn and unordered/α-helical structures^47–49^. This is to be expected given that β-sheet structures, being longer-range and generally rarer interactions, would likely emerge only in sequences longer than 25-mers or upon self-assembly of shorter peptides^50^. Combined, CD and IR spectroscopy therefore indicate that two alphabets beyond C20 can form familiar secondary structural elements.

To demonstrate that both xeno alphabets are capable of forming an α-helical structure under favorable conditions, we measured the CD spectra of the libraries after adding 2,2,2-trifluoroethanol (TFE), a cosolvent known for inducing α-helical structures. Following the addition of TFE, we indeed observed a change in the spectra, indicative of an increase in helical structure content for both Xeno libraries (see Extended Data Fig. 3a). While these results are consistent with small canonical peptide libraries, they serve as important controls showing that both Xeno alphabets bear intrinsic capacity to fold and do not present steric constraints that prevent formation of secondary structure arrangements, such as α-helices.

To further characterize the Xeno 1 and Xeno 2 combinatorial libraries, we investigated solubility (Extended Data Fig. 2), aggregation propensity, and secondary structure changes (Extended Data Fig. 3), across a range of pH values (pH 3, 5, 7.4, 9, 11). Overall, Xeno 2 emulates E10 in that it shows less solubility at lower pH (3 and 5) and more solubility at higher pH (7.4, 9, and 11). At lower pH however, Xeno 1 is more soluble than both Xeno 2 and E10. The changes in the CD spectra measured across the pH range (Extended Data Fig. 3b) most likely reflect this change in solubility (Extended Data Fig. 2). The difference in solubility may plausibly be explained by the (de)protonation of side chains of some amino acids (see Extended Data Table 1) and the various side chain polarities. Secondary structure propensity of Xeno 1 therefore appears to remain unchanged across the pH range. Size-exclusion chromatography (SEC) indicates no substantial oligomerization or assembly propensity in either xeno alphabet (Extended Data Fig. 3c). This contrasts with the substantial self-assembly propensity observed for E10^50^. Interestingly, neither solubility nor aggregation propensity were selected for directly during alphabet design. Solubility thus appears to result from selection for alphabets that emulate E10’s hydrophobicity range and distribution.

### Sequence design of xeno peptides and characterization of secondary structure

Because all results presented so far are based on combinatorial random sequence libraries, they demonstrate the *average* inherent propensity for Xeno 1 and Xeno 2 to form secondary structure. To assess the structure forming potential of Xeno 1 and Xeno 2 at a finer resolution, we therefore designed specific sequences with binary patterning theory^51^ to adopt α-helix, β-sheet, or a combination of the two with five motifs: anti-parallel adjacent α-helices connected by a loop (helix–loop**–**helix); two adjacent β-strands and α-helix connected by loops (ββα); and four adjacent β-strands connected by loops (See Extended Data Fig. 4).

For each of these five motifs we designed 20 sequences per alphabet, each comprising 25-26 amino acids (200 sequences in total listed in SI_Sequences.xlsx; see methods for details). Another set of 200 sequences was prepared as a control by random shuffling the designed sequences. From this pool of 400 candidates, 96 peptides were successfully synthesised by parallel solid-phase peptide synthesis in quantities sufficient for downstream analysis, of which 44 matched the target mass observed in LC-MS spectra. We measured electronic synchrotron radiation circular dichroism spectroscopy (SRCD) spectra of these 46 to exploit the high sensitivity of SRCD compared to standard CD measurements (see “SI_SRCD.xlsx” in the supplementary information). The spectra obtained suggest a varying average of secondary structure content. We therefore selected the 7 most promising peptides, in terms of secondary structure propensity (Fig. 4a), for more detailed inspection using temperature-dependent CD and NMR.

**Figure 4.**
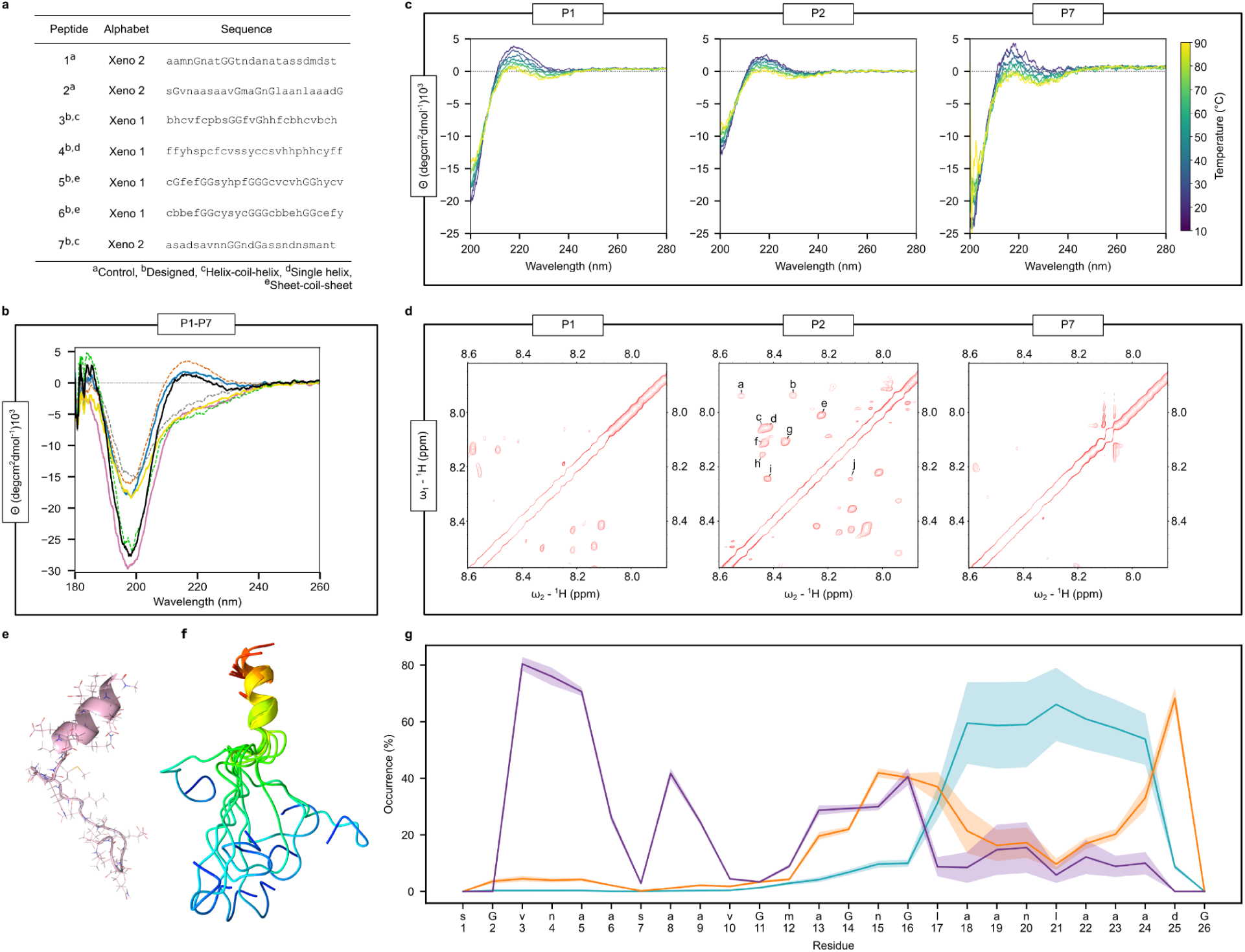
Secondary structure propensity characterization of individual xeno peptides via NMR and CD spectroscopy. **a,** Peptide sequences P1-P7 (using one-letter abbreviation described in Fig. 2) **b,** Electronic circular dichroism spectra for P1 (blue solid), P2 (orange dashed), P3 (pink solid) P4 (gray dashed), P5 (solid yellow), P6 (green dashed), and P7 (black solid). **c,** Temperature dependence of P1, P2, and P7. **d,** The amide regions (7.9–8.6 ppm) of P1, P2, and P7 in the 2D ^1^H–^1^H NOESY spectra. The amide region of P2 shows well-dispersed resonances and sequential NH–NH cross-peaks, consistent with local secondary structure (a: Nva3-Gly2, b: Nva3-Hgn4, c: Nle17-Gly16, d: Nle17-Gla18, e: Nle21-Gla23, f: Dmg25-Gla24, g: Dmg25-Gly26, h: Hgn15-Gly14, i: Gla19-Gla18, j: Gla19-Hgn20). In contrast, the spectra of P1 and P7 exhibit few amide cross-peaks, indicating minimal ordered structural features. **e,** Representative 3D model of P2 with the lowest AMBER/GB energy from NMR-informed molecular dynamics **f**, Superimposed ensemble of P2 MD structures aligned to residues a18-a24 (right; colored by sequence position). **g,** Per-residue occurrence of secondary structure in P2 throughout the last 1.6 µs among 10 replicas of NOE-restrained MD simulations. Lines show the average, light areas delineate standard deviations. Single-letter xeno amino-acid codes are given. β-bend is shown in violet, β-turn in orange, and α-helix in cyan. Structural motifs negligibly present within P2 (pi-helix, 3-10 helix, β-bridge, and β-sheet) are not shown (see Extended Data Fig. 7).

Control peptides 1-2 (P1-P2), and binary-designed peptides 3-7 (P3-P7) were characterized using CD spectroscopy (Fig. 4a-b). P3, P4, P5 and P6 exhibited a negative spectral band at ∼198 nm, varying in intensity, along with a negative shoulder around 220 nm. This combination of spectral features leaves ambiguity to the extent of PPII structure versus disordered content. Interestingly, CD spectra for P1, P2 and P7 were characterized by a negative spectral band at ∼198 nm, varying in intensity, along with a positive spectral band at ∼215 nm, with an intensity approximately five times smaller. This spectral shape could be attributed to a peptide with a PPII structure^46,47^, the presence of which was supported by temperature dependent CD spectra (Fig. 4c). However, the presence of β-turns cannot be excluded^43^. More broadly, the additional carboxyl groups of Gla, for example, and their possible chiral arrangement can make structure assignment ambiguous within this CD spectra: higher-resolution techniques are thus required to dissect the possible spectral features of this peptide.

Interestingly, P1, P2, and P7 all derive from the Xeno 2 alphabet, tempting an interpretation that Xeno 2 presents greater structure-bearing propensity than Xeno 1. However, observed CD spectra of combinatorial libraries (Fig. 3a) do not directly support this assumption and our results derive from a meagre sample of 400 sequences drawn from a sequence space of 2 × 10^25^. It is more accurate to conclude therefore that it remains unclear whether Xeno 1 or Xeno 2 form structure more readily.

Two-dimensional NMR spectroscopy was performed to investigate whether P1, P2, and P7 exhibit any tendency toward secondary structure formation or higher-order assembly in solution. For P1 and P7 , the spectra did not show solid evidence of defined stable secondary structure (Fig. 4d). The resonances were poorly dispersed and highly degenerate, which is consistent with the limited amino acid diversity in these sequences. This degeneracy hindered unambiguous resonance assignment and precluded further structural analysis. In contrast, P2 displayed well-dispersed resonances, suggesting a more ordered local environment (Fig. 4d). Using TOCSY and NOESY experiments, most spin systems could be assigned (Extended Data Fig. 5). The NOESY spectra showed clear signals in the Hα region (4.1 – 4.5 ppm; see Extended Data Fig. 5a), as well as cross-peaks characteristic of NH–NH correlations between consecutive residues in the amide region. Additional correlations were observed in the aliphatic region (Extended Data Fig. 5b), corresponding to β- and γ-protons resonances of side chains as well as resonances of side chain terminal methyl groups. The presence of sequential NOE cross-peaks between spatially neighboring residues further supports a defined local conformation in P2.

Using NMR-informed molecular dynamics, we have observed a propensity towards a short β-bend at the N-terminus (residues v3-a5) and an α-helical structure at the C-terminus (residues a18-a24; Fig 4e-f). The φ and ψ dihedral angles of these residues in 10 representative structures from independent MD replicas suggest a mix of C-terminal α-helix and N-terminal β-bend, except for MD replicate number 4, the least stable structure, where we observe a shorter C-terminal α-helix starting at n20 (Extended Data Fig. 6). P2 is thus the first peptide constructed entirely from a xeno amino acid alphabet that has been observed to adopt any secondary structure elements according to CD, MD, and NMR.

## Conclusions

In this work, we demonstrate that two sets of amino acids, each quite different from the set of 20 found within the genetic code, can be polymerized into peptides which exhibit familiar, fundamental secondary structure. This successful demonstration of alternative, structure-forming *sets*, or xeno alphabets (Fig. 2), advances a rapidly maturing biotechnological landscape of engineering *individual* xeno amino acids into otherwise “natural” peptides and proteins^52^. Specifically, our results open the door to a separate protein-fold universe with which to test where xeno protein structures emulate, and where they diverge from, all that has been learned from C20 about protein folding. The possibility that further biophysics awaits discovery certainly resonates with a recent systematic search of dipeptides built from chemically diverse xeno amino acids which identified novel structures unknown from natural proteins^26^. Xeno amino acid alphabets which demonstrate foldable peptide structures thus seem likely to advance the “Principles that Govern the Folding of Protein Chains.”^2^

Far less clear are the boundary conditions which define a “foldable set” of amino acids. To approach the uncharted and effectively infinite number of theoretical amino acid sets^30^, both xeno alphabets tested here were designed to differ significantly from, while maintaining subtle similarity to, the canonical alphabet C20. Dissimilarity comes from choosing different sidechains, even different heteroatoms; Similarity comes from projecting a simple, physicochemical profile of E10 onto the chemical space of possible candidates (Extended Data Fig. 1). The resulting evidence that xeno alphabets can be polymerized into structure-bearing peptides thus highlights the importance of asking: how difficult is it to define a structure-bearing set of amino acids? It is currently unknown whether future improvements to the design protocol used here (Fig. 1) will severely constrain the flexibility of structure-bearing alphabets or whether xeno alphabet design is limited only by the imagination of researchers.

In the meantime, the mere existence of structure-forming xeno alphabets is significant to structural biochemistry, biotechnology, and astrobiology. For structural biochemistry, xeno alphabets allow future research to test the extent to which science has successfully distilled, from C20, the fundamental physics by which peptides form three-dimensional structures. For biotechnology, evidence that xeno alphabets can bear structure advances the frontier of generating novel “*structural motifs for the construction of new materials and macromolecules*”^26^ because new functional groups and even new heteroatoms enable chemistries beyond anything possible with C20. Finally, from the perspective of astrobiology, empirical examples of structure-bearing xeno peptides inform how easily an independent origin of life might construct metabolism from a different set of molecular building blocks.

The shared challenge faced currently by all of these frontiers is the lack of scientific infrastructure compatible with amino acids beyond C20. Specifically, there are no tools for xeno peptide design, no optimised synthesis protocols to polymerize xeno peptides, and no commercial instrumentation to sequence any such peptides. Our results thus motivate the development of this infrastructure to uncover the full potential of xeno amino acids.

## Methods

### Alphabet Searching & Filtering

#### Defining options for an xeno amino acid alphabet

All analyses presented here explore the amino acids available as ɑ-N-Fmoc and sidechain-protected derivatives within the IRIS Biotech GmbH commercial supplier (https://www.iris-biotech.de). This resulted in a collection of 248 monosubstituted Fmoc-L-α-amino acids suitable for solid-phase peptide synthesis.

#### Selecting a set of xeno amino acids which resemble E10

Xeno amino acid *alphabets* were then identified within this collection using a heuristic search designed to emulate patterns of physicochemical coverage seen in the E10 amino acid alphabet (see SI.1). The fitness function guiding this search calculated four-dimensional euclidean distance abstracted from physicochemical coverage^28,53^ using van der Waals Volume (*calculated with KNIME VBAC*^54^ *ver. 4.3.0.v21000*) and ChemAxon JChem logD at pH 7.0 (*ver. 21.10.0,* https://www.chemaxon.com). See previous work^55^ protocol §5 for detailed instructions describing how to computationally calculate these descriptors. Fitness was then defined as the Euclidean distance in coverage from one xeno alphabet to E10 multiplied by the inverse sum of volumes in the xeno alphabet:

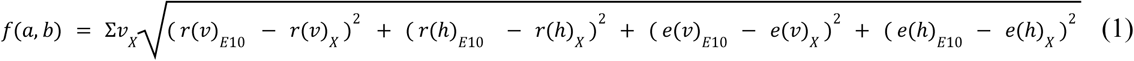

Where E10 is the early ten amino acid alphabet, *X* is the xeno alphabet being evaluated, *v* is the calculated van der Waals volume, *h* is the calculated hydrophobicity (LogD, pH 7.0), *e* is the evenness of each descriptor (*v, h*), *r* is the range of a given descriptor, and Σ*v_X_* is the sum of van der Waals volumes of all amino acids in a given xeno alphabet. The weight metric Σ*v_X_*ensures that alphabets mimic the specific range of volumes found within E10 (see also Extended Data Fig. 1).

The heuristic search begins with a random xeno amino acid alphabet comprising nine xAAs and glycine (Gly). After calculating the alphabet’s fitness, one of the nine xAAs is replaced and the fitness is recalculated. Unless the fitness improves, the change is rejected. This process repeats for 100 iterations (successful accepted substitutions) before reporting an optimised alphabet. The results of 10,000 searches of the IRIS Biotech library produced 1,044 unique, non-redundant xeno amino acid alphabets comprising 166 unique xAAs (because there are a subset of xAAs predisposed to form sets which emulate life’s alphabet)^28^. These 1,044 were then filtered to 26 which match the formal charge of E10 sidechains by comprising 2 negatively charged, and 8 neutral sidechains at pH 7.0 in water.

### Xeno Alphabet Selection

For all unique xAAs within the top 10 highest physicochemical coverage alphabets (SI_Xeno_Candidates.xlsx) as well as 20 canonical amino acids, we used the model of an amino acid with acetylated N-terminus and N-methylamidated C-terminus to better emulate their behaviour in proteins without artificial terminal charge. All amino acids were considered in their natural protonation state in water (Asp, Glu charge −1; His, Lys, Arg charge +1) in all calculations. Comparing estimated p*K*_A_ of the four negatively charged xeno amino acids, two per alphabet, with both the experimental^56^ (3.90 and 4.07 for Asp and Glu resp.) and calculated p*K*_A_ of Asp and Glu confirmed their deprotonation at pH 7.0 (see Extended Data Table 1). RDkit ver. 2023.9.4^57^ was used to generate the starting geometries (one for each xAA). To sample the whole potential energy surface (PES) of the amino acid backbone, each *φ* and *ψ* backbone dihedral angle was optimised with constraints to adopt a discrete value from the list {−180°, **−**150°, −120°, −90°, −60°, −30°, 0°, 30°, 60°, 90°, 120°, 150°}, resulting in (−180…150)⨯(−180…150) matrix of the PES with 144 possible backbone dihedral angle combinations. This preliminary optimization was carried out with UFF force field^58^ (implemented in RDkit), iteratively, starting with force constant 0.002 and sequentially increasing it until both *φ* and *ψ* angles were within 5° of the desired values. Based on our extensive previous experience^59–61^ in generating large sets of peptide conformers, we then used the CREST program (Conformer Rotamer Ensemble Sampling Tool, ver. 2.12)^62^ for sampling of the amino acid side chains, keeping the backbone *φ* and *ψ* dihedral angles constrained at target values, leaving the side chain dihedral angles for free sampling. Due to the well recognized problem of commonly used force fields to overestimate the stability of α-helix^63–67^, the GFN2-xTB semiempirical QM method^68^ was used for conformer optimization and the ALPB implicit solvation model^69^ was used for modelling the solvent effects, with water as solvent. As we consider the GFN2-xTB single point energies insufficiently accurate, we carried out the DFT single point energy calculation on the GFN2-xTB optimised conformers instead of the GFN2-xTB semiempirical single point, directly during the CREST conformer search. This DFT single point calculation was performed using TURBOMOLE (ver. 7.6)^70^. We employed the BP86 functional^71^, DGauss-DZVP basis set^72^ and Grimme’s D3(BJ) dispersion correction with special parameters for proteins^73,74^. Solvation effects for water as solvent were treated with the COSMO (conductor-like screening model)^75,76^ and COSMO-RS (COSMO for realistic solvation)^77,78^ solvation models as implemented in the BIOVIA COSMOtherm (ver. 23) program. The “BP_TZVPD_FINE_23.ctd” parametrization file with FINE cavities^79^ was used. Final free energies of conformers (*G*) were calculated within COSMOtherm program, according to the following formula:

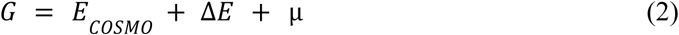

Where *E*_COSMO_ is BP86-D3BJ/COSMO (*ε* = ∞) energy of the conformer, Δ*E* is the averaged correction for the dielectric energy, and *μ* is the chemical potential of the conformer. The integration from an ideal conductor to the real solvent with a given permittivity is included in the Δ*E* and *μ* terms, and is done inherently within the COSMO-RS procedure. All three values are summed in the COSMOtherm program. The structure with the lowest DFT(BP86-D3)/DGauss-DZVP//COSMO-RS energy was then considered as a representative conformer for the given *φ*,*ψ* combination on the PES.

For one *φ*,*ψ* combination of 3,3-dimethylproline (150°,−150°), no conformers were successfully optimised and therefore were not calculated. This missing conformer belongs to the high energy region of the ring bending of this proline-like amino acid (*φ* > 90°), therefore we decided to ignore it as unimportant. In addition to systematic conformational searches carried out for the capped xeno amino acids, we conducted conformational sampling and DFT calculations for all possible capped xeno tripeptides formed by xeno amino acids within each alphabet (see SI_PES.html).

#### Ramachandran distance metrics

In order to assess the ‘similarity’ of canonical and xeno amino acid alphabets in some rigorous way, we wanted to compare these alphabets in terms of PES. As direct comparison of PES would be difficult, we used the Principal Component Analysis (PCA)^80^ as implemented in SciPy (ver. 1.12.0)^81^ to represent each PES as a point in 2D space (SI_PCA.html). The PCA transformation was performed on the whole PES (12⨯12 matrix of representative conformers) as follows: for all representative conformers, the Boltzmann probability was calculated. Conformational energies higher than 5 kcal·mol^−1^ were set equal to 5 kcal·mol^−1^ (missing conformer of 3,3-dimethylproline was treated in the same way, as it represents high energy region on PES). Subsequently, we constructed two principal components out of Boltzmann factors of each point of PES for 20 canonical amino acids, and then used these components to represent all xAAs in the same 2D space (see Extended Data Fig. 1).

Using PCA results, we used linear sum assignment (LSA) analysis to assess which of the final candidate alphabets are closest to the E10 alphabet in terms of Ramachandran plots of specific amino acids. LSA gives a one-to-one mapping of each amino acid in one alphabet to one amino acid in another (reference) alphabet. Using this one-to-one mapping, we compared each of the ten xeno alphabets to E10 and ranked them by how similar they are to E10. To find the mappings, for each alphabet, we calculated the Euclidean distance between the position of every amino acid in a given alphabet in PCA and the position of every amino acid in E10 in PCA (See Extended Data Fig. 1c-d) using *cdist* from SciPy (ver. 1.12.0)^81^. LSA then finds such one-to-one mapping, where the sum of these distances is the lowest. All considered alphabets were then ranked according to this distance. This was calculated using the linear_sum_assignment function as implemented in SciPy^82^. One of the reasons we wanted to use LSA as a criterion is that if we want to emulate natural folds with xeno alphabets, LSA allows us to find the amino acid with the closest Ramachandran plot to an amino acid in the natural fold, thus raising the chance that the fold will prefer the same (or very similar) conformations.

We additionally used Earth Mover’s Distance (EMD), also known as Wasserstein Distance, as another metric to calculate the differences of each alphabet’s PES (see above) to E10. EMD is a similarity metric used when comparing two distributions. EMD calculates the amount of ‘work’ required to move one distribution to match another^83^. In order to calculate the ‘work’ required to transform one PES to another, we first flattened each amino acid’s PES into a one-dimensional array. Given that each position in this array represents a specific pair of *φ*,*ψ* torsion angles and that the value at each position represents the conformational energy, we could then calculate the Wasserstein Distance from each xeno amino acid in a given alphabet to those within E10 using the *wasserstein_distance* function as implemented in SciPy. Finally, we report each alphabet’s mean Wasserstein Distance from E10 as a quantitative measure of the distance between a xeno alphabet’s PES and E10’s PES (see Extended Data Table 2).

Using these metrics, we selected two final xeno amino acid alphabets to further characterize experimentally. The final decision was based not only on the PCA/LSA and EMD scoring, but also on the presumed stability and predicted ability to polymerize to guide design of xeno peptides (see below).

#### Linear Sum Assignment (LSA) & Earth Movers Distance (EMD)

In Extended Data Table 2, LSA values represent the sum of the distances in the best mapping of a given alphabet onto E10 within PCA coordinates, and EMD values represent the mean Wasserstein Distance from the PES for each amino acid in the respective alphabet to each amino acid within the E10 alphabet.

#### HOMO-LUMO gaps

As an additional characteristic of the xAAs, we calculated the HOMO-LUMO gaps for both Xeno 1 and Xeno 2 alphabets and also for 20 canonical AAs (early 10 + late 10). For each amino acid, the conformer with lowest energy in constrained sampling (see above) was subsequently optimised without constraints, on the DFT(BP86-D3BJ)/DGauss-DZVP//COSMO in water (*ε_r_*=80) as solvent. HOMO-LUMO gaps were then calculated on the optimised geometries at HF^84^/def2-TZVPD//COSMO(*ε_r_*=80) level. The calculations were performed using the same program package and all other settings as seen in the energy calculations.

#### Theoretical estimation of solubility of xeno amino acids

To estimate the solubility in water, we used the ESOL log(S) model^85^ as implemented at SwissADME web tool^86^. The estimated solubilities of 20 canonical amino acids obtained from this model correlate the best (R^2^=0.79, see SI_Solubility_Benchmark.xlsx in the SI), out of models tested in our benchmark, with the established Wimley-White whole-residue hydrophobicity interface (water to bilayer) scale^87^. While solubility in water and hydrophobicity are two different quantities, these are closely related and thus ESOL log(S) should, in principle, be a quite viable model for the solubility estimation for the xeno amino acids. We note that the model for the solubility estimation is, expectedly, not perfect: The predicted and solubility of Gly and Ccs are too high and low, respectively, considering the bare amino acid backbone of Gly and polarity of the carboxylate in Ccs.

#### Sequence design

For the *ab initio* design of candidate xeno peptides with defined secondary structure, there is unfortunately no viable tool in contemporary (bio)chemistry^88^. Within the last year, multiple machine-learning based methods have emerged (in preprint) for design of (usually cyclic) peptides including non-canonical amino acids, but not for the predictions of shape and structure of linear fully xeno peptides^89–91^. Therefore, we employed the binary patterning theory^51^ using simple polar/non-polar categories for amino acids. According to the theory^51^, the helical and extended motifs are dictated mainly by the right combination of the hydrophobicity of the amino acid side chains (see Extended Data Fig. 3 and Extended Data Table 3). For the design, we consider Ccs (c), Pho (f), Oms (s), Omh (h) as polar, and Dmp (p), Pyr (y), Nva (v) as non-polar concerning the Xeno1 alphabet. For the Xeno2 alphabet, we consider Gla (a), Hgn (n), Oms (s) as polar and Mse (m), Nle (l), Nva (v), Dmp (d) as non-polar. The other amino acids, namely Gly (G), Abu, (b) and Mth (t), we denote as ‘other’. These assignments were made roughly according to the ESOL log(S) solubilities, with the exception of Ccs, which bears a negatively charged group but is predicted to be non-polar. We considered 5 secondary structure motifs, widespread in the existing proteins^92^: single α-helix, single β-strand, two anti-parallel adjacent α-helices connected by a loop (helix-loop-helix motif), two adjacent β-strands and α-helix connected by loops (ββα motif), and four adjacent β-strands connected by loops (Extended Data Fig. 3). As neither Xeno alphabet has positively charged amino acids, the designed sequences cannot benefit from the attractive +/− interactions, which is not a major disadvantage for the peptides comprising canonical amino acids.^93^ For each of the 5 motifs, we designed 10+10 sequences as follows: for the first 10 sequences, we randomly choose one AA from the list of polar or non-polar xAAs based on the template for the respective motif pattern, with 5% chance for choosing one of the unclassified xAAs instead, ignoring the pattern. Loops were composed mainly of Gly, to enhance the flexibility. The second 10 sequences were generated in the same way as the first 10 sequences, but were then ‘augmented’ with 1-5 triplets that showed propensity for particular secondary structure as isolated tripeptides in the calculations (see SI.2), keeping the binary patterning criterion as described. This resulted in 100 designed xeno peptide sequences (50 with pure random binary patterning and 50 with use of the isolated tripeptide preferences) for each xeno alphabet and 100 control sequences (200 sequences in total).

### Experimental

#### Synthesis of pooled libraries

The following procedure was used to produce both presented alphabets. 83 mg of Rink Amide AM resin (0.64 mmol/g) was swelled in 1 ml of DMF for 30 min, and deprotected. This was done by first washing the resin with 0.5 ml of 20% (v/v) piperidine in DMF on a centrifugal filter, and then incubating with additional 0.5 ml piperidine solution for 30 min. Resin was washed on a centrifugal filter unit (Biorad 7326204) 8 times with 0.5 ml of DMF and split into 10 equal portions as a DMF slurry into 1.5 ml PP tubes, and the residual DMF was drained with a pipet tip. To each portion, 70 µl (residues 1-10) or 140 µl (residues 11-25) of 300 mM solution of one of the ten AAs was added, followed by 30 µl or 60 µl of 875 mM solution of Oxyma Pure, and 4 µl or 8 µl of N,N′-diisopropylcarbodiimide. Couplings then proceeded o/n at RT on a carousel at 20 RPM, protected from light. The following day, all the split reactions were pooled, washed on a filter unit 4 times with 0.5 ml of DMF, and the next cycle was commenced by deprotection with piperidine as described above.

After final Fmoc cleavage, the resin was washed 8 times with DCM, and dried by spinning at 1000 × g for 10 min. Resin was then treated with 2 ml of 95:2.5:2.5 TFA:TIS:water mix for 2 h on a carousel at 20 RPM, protected from light. Resin was removed by filtration, and collected filtrate was precipitated with diethyl ether (DEE) by transferring the filtrate in equal portions into 14 ml of DEE in three 15 ml conical tubes. These were mixed briefly by inverting and submerged in liquid N_2_ until the solvent froze. Tubes were then spun for 5250 × g for 20 min in a centrifuge pre-chilled at 4 °C. Supernatant was removed by slowly inverting the tubes upside down and allowing the inverted tubes to drain on a paper towel. In each tube, the precipitate was resuspended in 5 ml of DEE, frozen in liquid N_2_ and spun again under the same conditions. Washing was repeated for a total of 3 times. Pellets were then allowed to dry in the open for 10 min, and then were resuspended and collected into a single tube in 5 ml of 1 mM HCl. Suspension was then lyophilized o/n, and the following day the material was resuspended in 5 ml of 1 mM HCl, frozen in liquid N_2_ and lyophilized again for a total of 4 times.

#### Individual peptide synthesis for structural validation

Peptides were synthesized on Rink Amide AM resin (0.61 mmol/g) by standard Fmoc SPPS using the automated parallel peptide synthesizer SPENSER (IOCB Prague). Coupling was performed with 300 mM Fmoc-protected amino acids and 375 mM Oxyma Pure in DMF, activated with 5.5% (v/v) DIC (1.25 eq. relative to the amino acid). Global deprotection and cleavage from the resin were carried out using a mixture of trifluoroacetic acid (TFA, 88% v/v), triisopropylsilane (TIS, 2% v/v), DL-dithiothreitol (5% w/v), and water (5% w/w). Crude peptides were precipitated with cold DEE (20 ml DEE/1 ml TFA mixture) and purified by reversed-phase HPLC using a 10–60% acetonitrile gradient with 10 mM triethyl ammonium acetate as additive over 60 min. Pooled fractions were then lyophilized o/n, and the following day the material was resuspended in 1 mM HCl, frozen in liquid N_2_ and lyophilized again for a total of 4 times.

#### General procedures

Commercially available HPLC-grade ACN, DMF, DIC and building blocks were used without further purification. Chemicals were obtained from Sigma-Aldrich, Lach-Ner, Novabiochem, and Iris Biotech.

High-resolution ESI mass spectra were obtained on an LTQ Orbitrap XL hybrid FT mass spectrometer (Thermo Fisher Scientific). Ionization conditions in the ESI Orbitrap source were optimized as follows: sheath gas flow rate of 35 au, auxiliary gas flow rate of 10 au (N_2_), source voltage of 4.3 kV, capillary voltage of 40 V, capillary temperature of 275 °C, and tube lens voltage of 155 V. Analytical LC-MS was performed on an Agilent 1290 Infinity II Autoscale system (1.5 ml min^−1^ flow rate, gradient elution from 2 to 70% ACN over 30 min, 10 mM triethyl ammonium acetate as additive) with ELSD detection using a Phenomenex Luna Omega C18 analytical column (5 μm, 250 x 4.6 mm) to verify compound purity. All final compounds achieved a minimum purity of 95% with the exception of peptide 4 (94.69%). See SI.4 and “SI_Peptide_1-7_Synthesis” in SI for the synthesis data.

#### Solubility and aggregation analysis

Solubility and aggregation analyses were performed in universal Britton-Robinson (ABP) buffers consisting of 20 mM acetic acid, 20 mM phosphoric acid and 20 mM boric acid. The pH was adjusted with 5 M NaOH to 3.0, 5.0, 7.4, 9.0, or 11.0, and the ionic strength was adjusted with NaCl to conductivity that corresponds to the conductivity of 50 mM or 500 mM NaCl. Prior to analysis, stock solutions of peptide libraries were prepared by resuspending them in autoclaved MilliQ water to the final concentration of 5 mg·ml^−1^. Stock solutions were diluted into the corresponding ABP buffer to the final concentration of 0.5 mg·ml^−1^ and incubated for 30 min at room temperature with mild shaking (500 rpm). After incubation, the insoluble part was separated by centrifugation at 21,300 *g* for 30 min at 4 ℃, and the supernatant (cleared sample) was subsequently used for solubility and aggregation analyses.

The relative solubility of peptide libraries was estimated by UV-Vis absorption spectroscopy and fluorescence spectroscopy. UV-Vis absorption spectroscopy analysis was performed by measuring absorption at 215 nm in cleared samples on Nanodrop 2000 (Thermo Fisher Scientific). For fluorescence spectroscopy analysis, the relative solubility was estimated using a fluorescamine assay in a 96-well plate format. 90 μl of the cleared sample was mixed with 30 μl of 3 mg·ml^−1^ fluorescamine in DMSO, and the resulting mixture was incubated for 30 min in the dark at room temperature. The fluorescence was then recorded using the following parameters: excitation at 365 nm, emission at 470 nm. 0.5 mg·ml^−1^ solution of peptide libraries in DMSO were used as a standard.

The aggregation behaviour of peptide libraries was estimated by analytical size-exclusion chromatography. A 100 μl aliquot of cleared sample was loaded onto a Superdex 75 Increase 10/300 GL column (Cytiva) that was pre-equilibrated with two bed volumes of the corresponding 20 mM ABP buffer. The peptides were eluted from the column isocratically by one bed volume of the ABP buffer at 0.5 ml·min^−1^ flow rate at room temperature, and the elution of peptides was monitored by absorption at 215 nm.

#### Circular Dichroism Spectroscopy

The ECD spectra were measured using the Jasco 1500 spectropolarimeter equipped with the Peltier holder PTC-517. Temperature dependence of CD spectra was measured in the 200-280 nm spectral range from 10 to 90 °C with the step of 10 °C in a 0.5 mm path length quartz cell (scanning speed of 10 nm/min, response time of 8 seconds, scanning step 0.5 nm, 1 accumulation) at nominal concentration 0.2 mg/ml. After baseline correction, the final spectra were expressed as molar ellipticity (*q*) (deg.cm^2^.dmol^−1^) per residue.

The synchrotron radiation circular dichroism (SRCD) spectra of individual xeno peptides were recorded on the AU-CD beam line at the synchrotron light source ASTRID2 (Aarhus, Denmark). Crude xeno peptides were dissolved in 10 mM phosphate buffer (pH 7.4) to the nominal concentration of 1 mg/ml, incubated for 15 min at room temperature, and briefly spun down at 14000x *g* to remove the insoluble portion. 30 μl of supernatant was subsequently placed into a quartz cuvette with 0.1 or 0.2 mm path length (Hellma type 121), and the SRCD spectra were recorded in the 170−280 nm wavelength range, in 1 nm steps with a dwell time of 2s/pt, in triplicate. The three scans were averaged and a reference baseline spectrum subtracted. The measurements were repeated on a duplicate sample, and the CD signal (mdeg) was averaged. After the analysis, the precise concentration of xeno peptides in samples was determined by HPLC amino acid analysis (based on the amount of Gly) and used to convert the averaged CD signal (mdeg) to molar ellipticity per residue (deg⋅cm^2^⋅dmol^−1^). The data were deposited in Supplementary Information, see SI.5 and “SI_SRCD.xlsx” file.

#### Infrared spectroscopy

IR spectra were measured on the Nicolet™ iS50 FTIR spectrometer (Thermo Fisher Scientific, Waltham, MA, USA) using a standard mid-IR source, KBr beamsplitter and a nitrogen-cooled MCT detector (2 cm−1 spectral resolution, Happ–Genzel apodization function, 1000 scans) in the 4000–600 cm−1 spectral range. The cell compartment was purged by dry nitrogen during all the measurements. The samples were dissolved in a 50 mM PBS buffer (5 mg/ml nominal concentration). The CaF_2_ BioCell with a 8 μm path length (BioTools, Inc.) was used for the FTIR spectra measurements. Spectral data treatment (GRAMS/AI) included solvent scan subtraction and background correction (using a linear function). Final IR spectra were normalized to the amide I intensity maxima. Second derivatives of the IR spectra were calculated using the *Savitzky*-*Golay* numerical *algorithm*.

#### NMR Spectroscopy

NMR spectra were acquired at 25 °C on an 850 MHz Bruker Avance spectrometer equipped with a triple-resonance (15 N/ 13 C/ 1 H) cryoprobe. The samples were dissolved at 0.7-1.5 mM in a volume of 0.35 mL, in buffer (10 mM PBS), 5% D_2_O, 95% H_2_O. Two dimensional homonuclear spectra were measured (TOCSY – 70 ms mixing time and NOESY – 120 ms). All pulse programs were from the standard Bruker library. Spectra were processed in Bruker Topspin I and analyzed in NMRFAM-SPARKY^94^.

#### NMR-Informed Molecular Dynamics Structure Elucidation

*Parametrisation.* To be able to run molecular dynamics (MD) simulations with NOE restraints from NMR, the P2 xeno amino acids had to be parametrised consistently with the ff19SB^95^ and ff19SB_mod AMBER force fields used here for Gly and Mse, respectively. The individual xeno amino acids protonated according to pH=7.4 were capped by formyl and N-methylamide at the N- and C-termini and optimized using GFN2-xTB^68^ with fixed dihedrals to avoid formation of intramolecular H-bonds. The partial charges were obtained with RESP fitting^96^ of ESP potential at the HF/6-31G* level. The caps were removed and their charge was dispersed over all the atoms.

##### MD Setup

Using the LEaP program of AmberTools24 suite^97^, the fully extended P2 peptide capped by acetyl and N-methylamide at the N- and C-termini was assigned mbondi2 radii consistent with IGB=5 generalized Born implicit solvation model used here for MD^98^. Implicit ion strength of 0.15 M and infinite cutoff for non-bonded interactions were used. Using the sander program of AmberTools24 suite^97^, minimisation of 5000 cycles was followed by stepwise warming from 5 to 300 K and increasing the NOE restraints weight over 120 ps in NVT ensemble. The timestep of 1 fs was used. The Langevin thermostat with friction coefficient of 1 ps^−1^ was used. Thereafter, the pmemd.cuda program of Amber24 suite^97^ was used to run 1.9 µs-long production runs where the timestep was increased to 2 fs. MD was run in 10 independent replicas. Most NOE restraints were satisfied during MD; if we consider a distance violation of up to 0.1 Å, there were only 8 out of 164 which occurred in more than 10% of snapshots; no violation greater than 0.2 Å was observed.

#### MD Analyses

From the production MD (Extended Data Fig. 7), it appears that the first 300 ns were needed to achieve stable secondary structures. The first 300 ns of trajectories were therefore omitted from subsequent analysis. The φ and ψ dihedral angles over the last 1.6 μs were averaged for each trajectory so as to find one structure from MD with the highest similarity to the NMR structure. These 10 selected structures underwent 5000 cycles of unrestrained MM/GB5 minimisation to yield the final 10 representative structures. All the analyses were conducted using the cpptraj program of AmberTools24 suite^97^.

## Data Availability Statement

All data needed to evaluate the conclusions in the paper are present in the paper and/or the Supplementary Information.

## Acknowledgements

This work was supported by a research grant from the Human Frontier Science Program https://doi.org/10.52044/HFSP.RGEC272023.pc.gr.168579, the Ministry of Education, Youth and Sports of the Czech Republic through the e-INFRA CZ (ID:90254), the Czech Science Foundation (GA CR, grant 26-22168S to L.R.), and by the Charles University, project GA UK No. 164524 (T.K.). Some computational work was supported by using the *taki* cluster of the University of Maryland Baltimore County High Performance Computing Facility (HPCF). This project has received funding from the European Union’s Horizon 2020 research and innovation programme under grant agreement No 101004806 (MOSBRI-2024-277 project). The authors wish to thank Dr. Søren Vrønning Hoffmann and Dr. Nykola Jones for their expertise in SRCD analysis.

## Author Contributions

Conceptualization: S.M.B., T.K., R.K., L.R., S.F., K.H. ; Chemical Synthesis: R.K., V.Ver. ; Experimental Investigation: R.K., V.Ver., P.S., M.M., L.B., M.P, R.H. ; Computational Investigation: S.M.B., T.K., J.M.K., M.L., J.Ř., E.A.; Visualisation: S.M.B., T.K., J.M.K. ; Writing–original draft: S.M.B., T.K., R.K., J.M.K., M.M., E.A., S.F., K.H. ; Writing–editing: S.M.B., T.K., R.K., J.K., J.M.K., V.Ver., L.B., M.P., R.H., M.L., J.Ř., P.S., L.R., S.F., K.H., V.Vev. ; Supervision: S.F., K.H., L.R., V.Vev. ; Project Administration: S.F., K.H. ; Funding Acquisition: T.K., S.F., K.H., L.R.

## Additional information

**Supplementary Information** is available for this paper.

**Correspondence and requests for materials** should be addressed to Drs. Sean M. Brown, Tadeas Kalvoda, Klára Hlouchová, and Stephen Freeland.

## Extended Data Figures

**Extended Data Figure 1.**
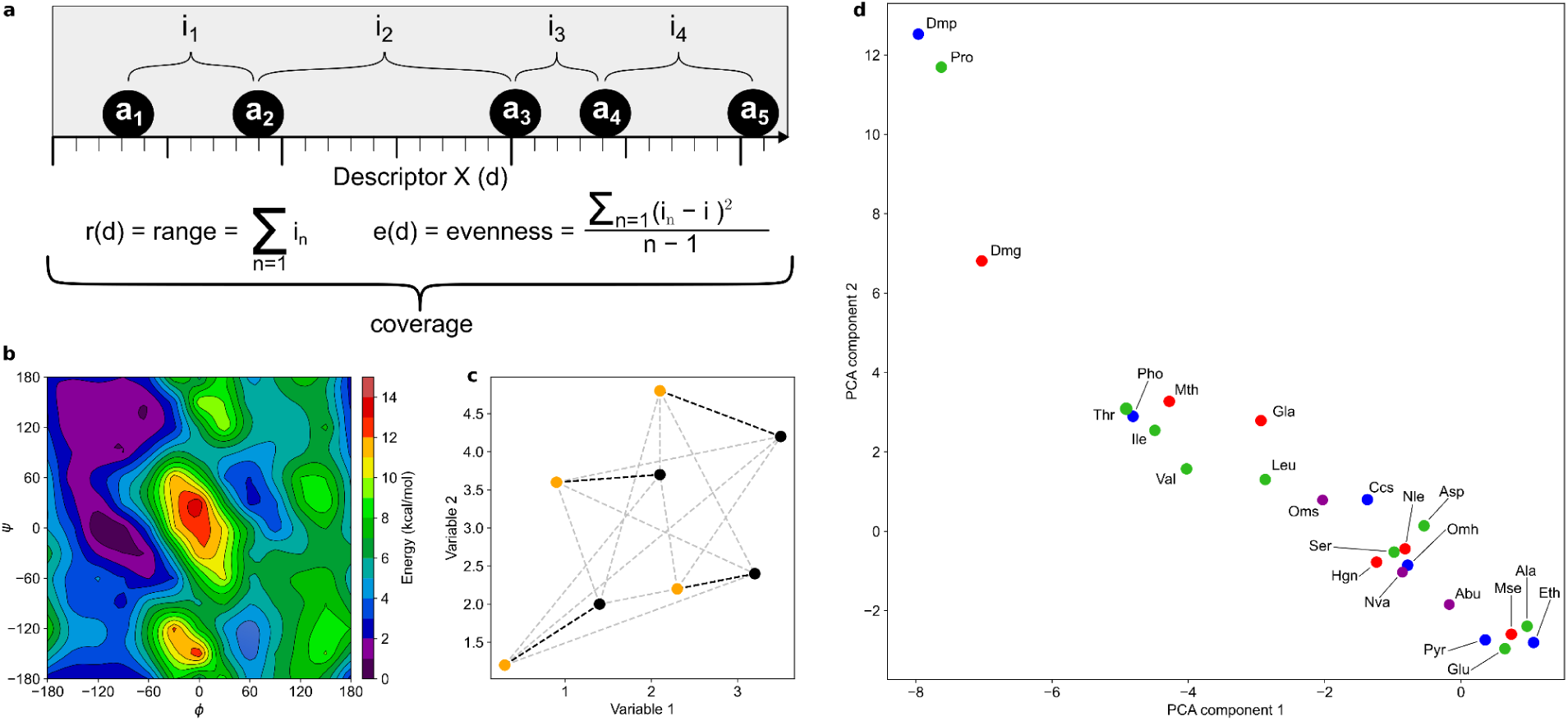
Xeno alphabet selection. **a,** Definition of Range and Evenness. For a given chemical descriptor such as van der Waals volume, “coverage” combines two statistics to represent a set of amino acids. This figure illustrates these statistics for a set of five amino acids (*a_1_…a_5_*) with four corresponding intervals (*i_1_…i_4_*) measured in terms of the hypothetical quantitative ‘descriptor x’ (*d*), for example van der Waals Volume. Evenness (*e*) is the sample variance of intervals between amino acids (*i*); Range (*r*) is the sum of these intervals (∑*i_1…4_*); “Coverage” is therefore the combination of range AND evenness for any given physicochemical descriptor. **b,** An example of an analysed potential energy surface for amino acid Ccy. **c,** LSA Scheme. For a given set of points (gold), which are to be mapped onto another set of points (black), the distance matrix is calculated between the two sets (grey dashed lines). LSA finds the one-to-one mapping which minimises the sum of these distances (black dashed lines). **d,** The actual PCA components of E10 (green), Xeno 1 (blue), and Xeno 2 (red) alphabets. Amino acids shared by both alphabets are depicted in magenta. Glycine (coordinates 19.8, 7.6) is omitted from the plot as it would always be assigned to itself.

**Extended Data Figure 2.**
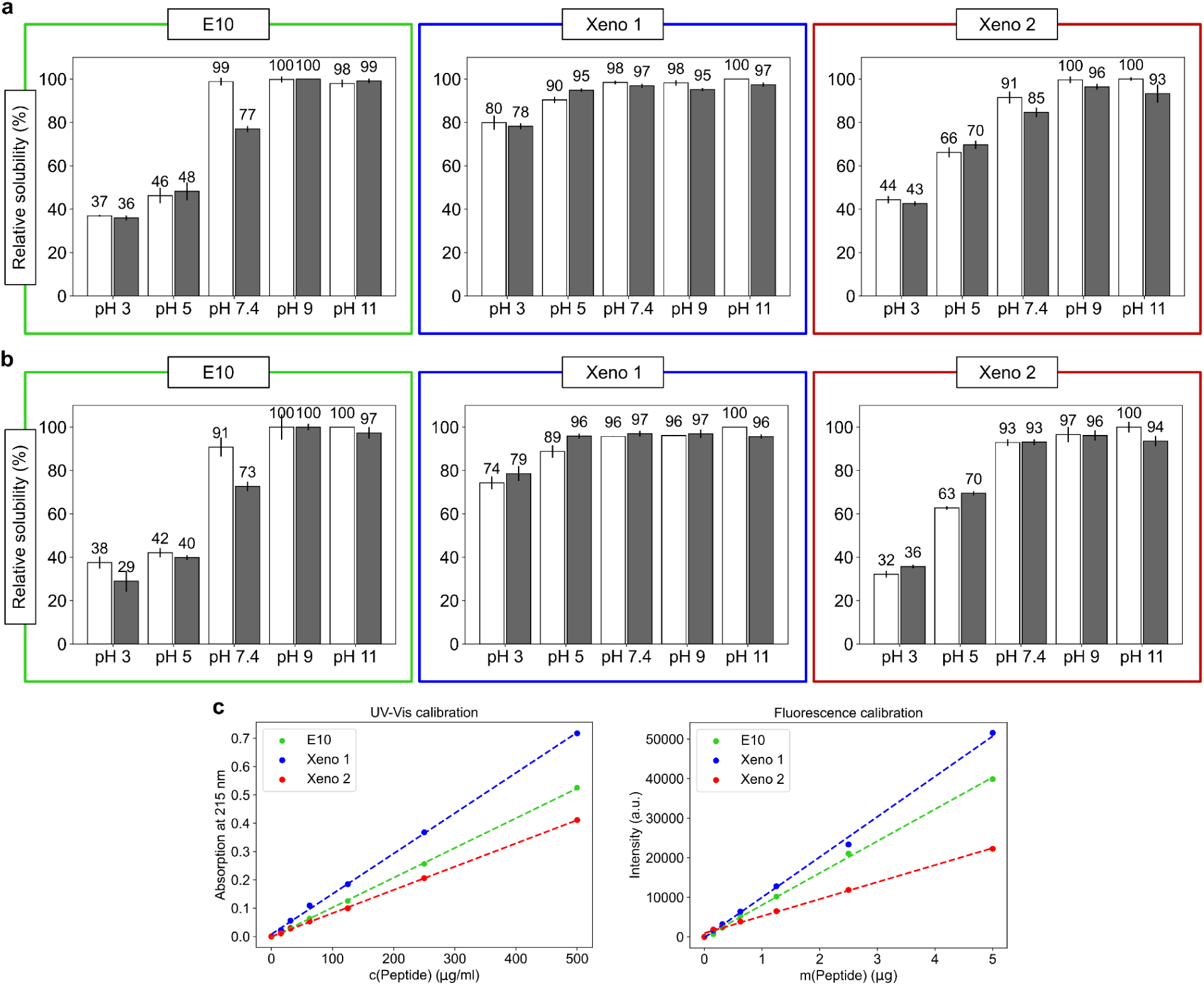
Solubility of the E10, Xeno 1, and Xeno 2 alphabets. **a,** Relative Solubility of 25mer libraries measured by UV-Vis Absorption Spectroscopy. White and grey bars correspond to medium (50 mM NaCl) and high (500 mM NaCl) ionic strength, respectively. Error bars are standard deviations between duplicates. E10 data was adapted from previous work^15^, as stated in methods. **b,** Relative Solubility of 25mer libraries measured by Fluorescence Spectroscopy. White and grey bars correspond to medium (50 mM NaCl) and high (500 mM NaCl) ionic strength, respectively. Error bars are standard deviations between duplicates. E10 data was taken from the same previous work^15^, as stated in methods. **c**, Solubility calibration curves for 25mer libraries for UV-Vis absorption (top) and fluorescence (bottom) spectroscopy: (i) E10 light green, (ii) Xeno 1 - blue, (iii) Xeno 2 - red.

**Extended Data Figure 3.**
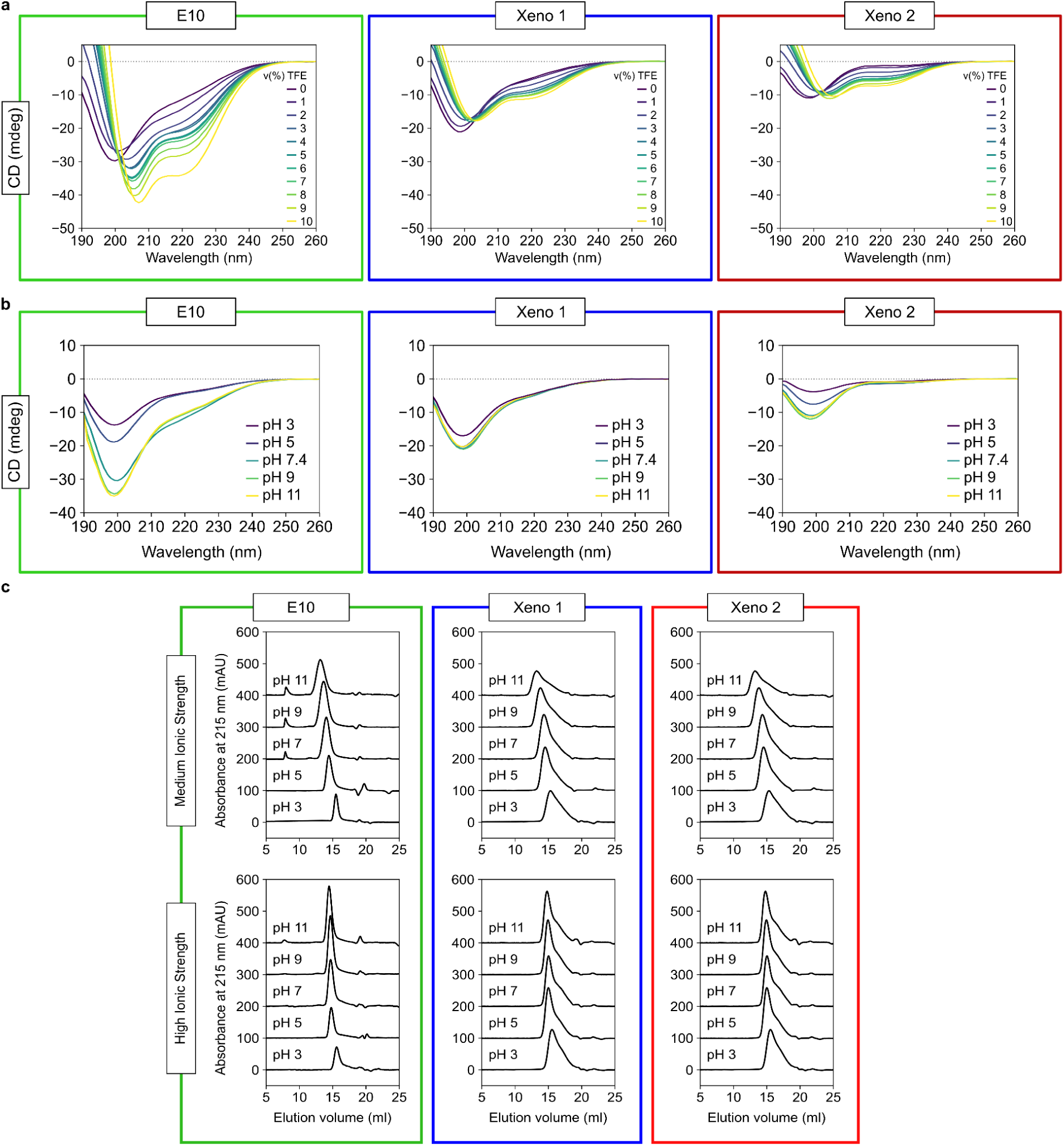
Additional characteristics of the E10, Xeno 1, and Xeno 2 alphabets. **a,** Induction of secondary structure with 2,2,2-trifluoroethanol (TFE). Spectra shown were measured at 0.2 mg/mL nominal concentrations of the peptide libraries in 10 mM ABP buffer (pH 7.4). See SI.3 for methods. **b,** Far-UV circular dichroism (CD) spectra at five pH values. The spectra of E10, Xeno 1, and Xeno 2 are representative of duplicates. See SI.3 for methods. Data for E10 was taken from previous work.^25^ The spectra were measured at 0.2 mg/mL nominal concentrations of the peptide libraries. **c,** aggregation propensity of E10, Xeno 1, and Xeno 2. Aggregation propensity was measured in 20 mM ABP buffer at various pH’s and medium (50 mM NaCl) and high (50 mM NaCl) ionic strength with size-exclusion chromatography (SEC). SEC chromatograms report absorbance at 215 nm as a function of elution volume. Respective chromatograms were shifted by 100 mAU for comparison.

**Extended Data Figure 4.**
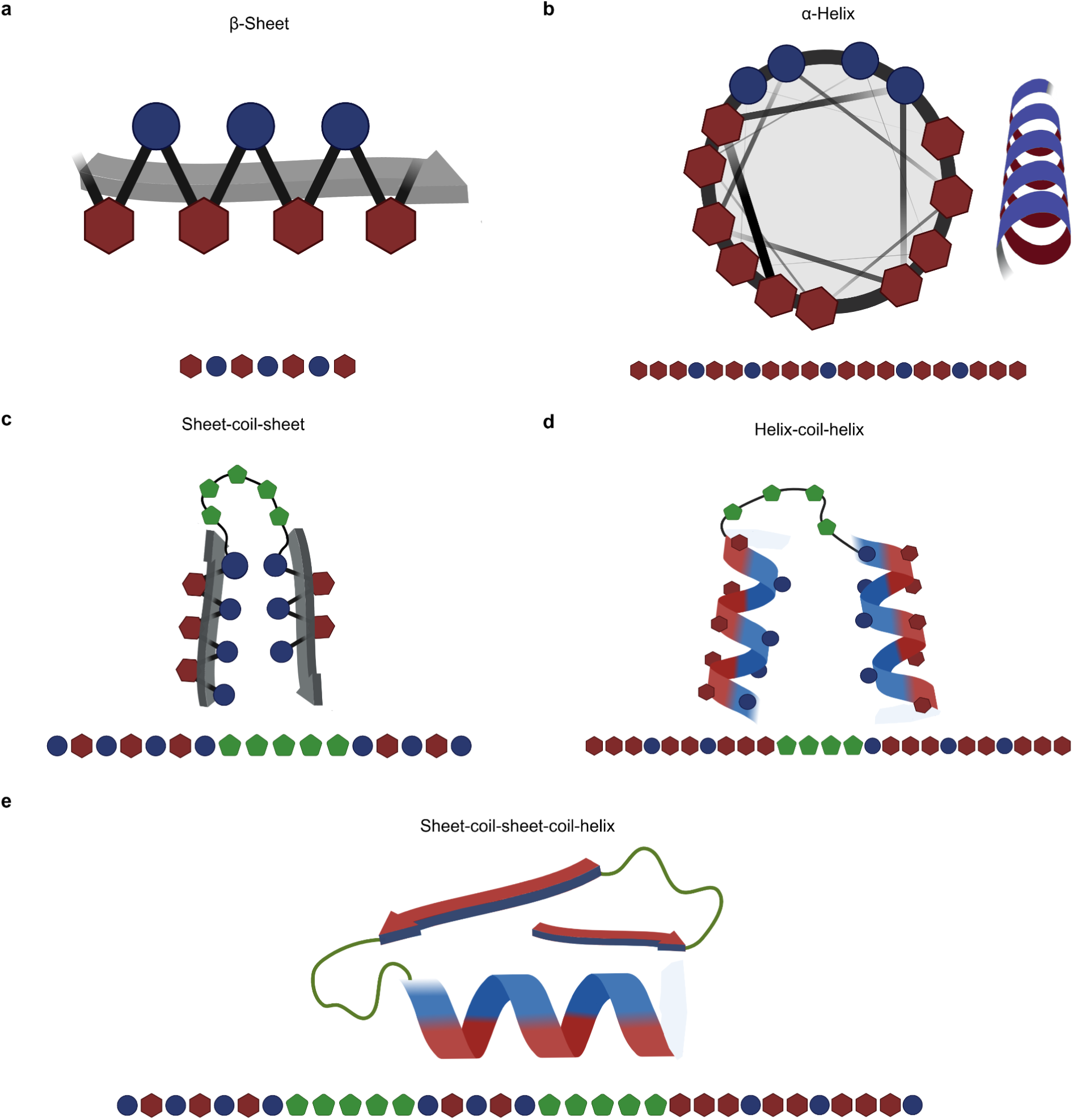
Binary Patterning Theory used for xeno peptide design. **a**, single β-sheet, **b**, single α-helix, **c,** sheet-coil-sheet (part of the four adjacent β-strands connected by loops, used in the design), **d,** helix-coil-helix. **e,** the motif of two adjacent β-strands and α-helix connected by loops (ββα motif). Polar residues shown in red hexagons, non-polar residues in blue circles, and other residues (e.g. Gly) in green pentagons. For the specific sequence patterns used in this study, see Table S1. Created in BioRender. Brown, S. (2026) https://BioRender.com/k98n286

**Extended Data Figure 5.**
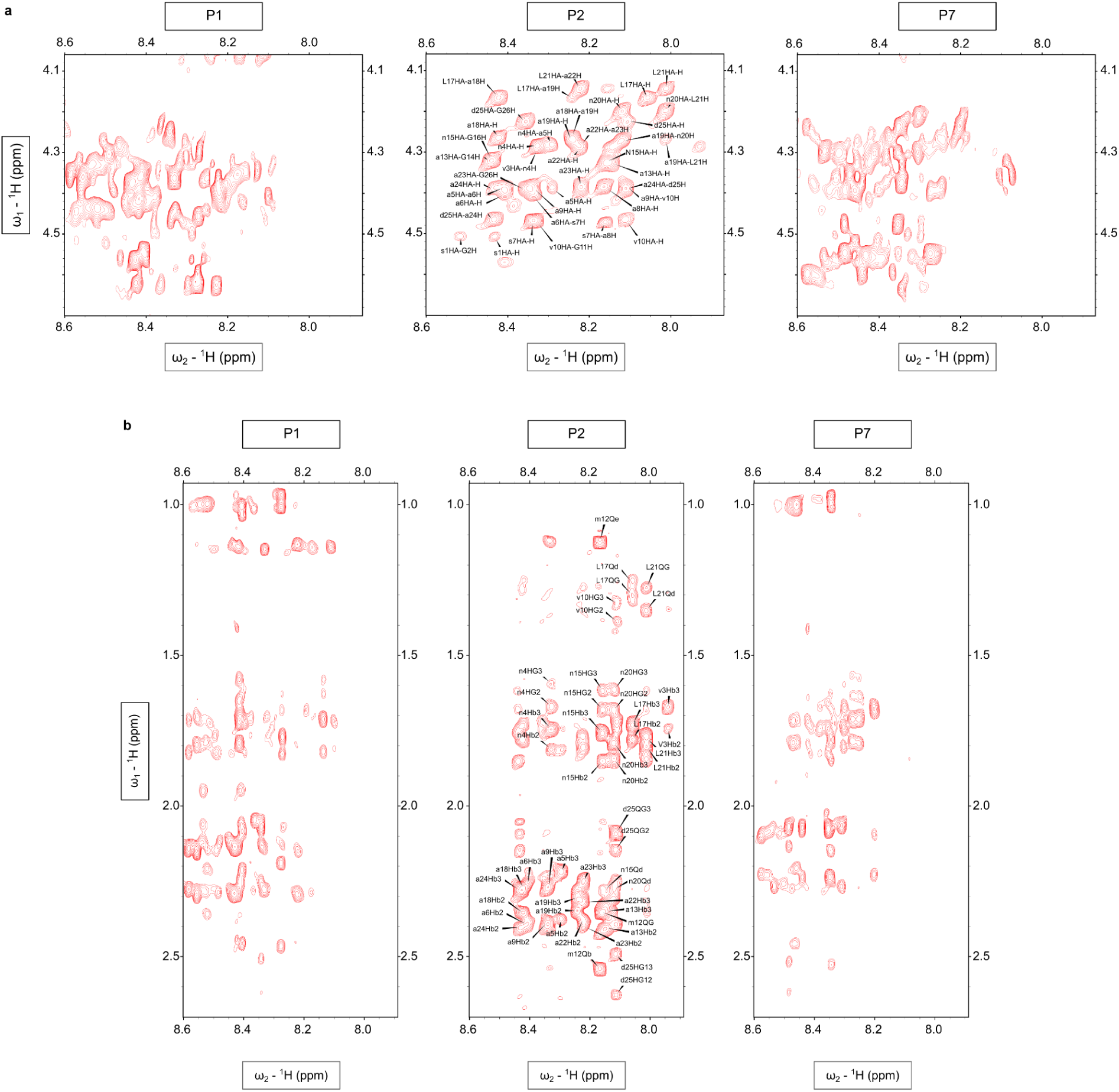
Additional NMR data of inspected peptides. **a**, Comparison of the Hα regions (4.1–4.6 ppm) of P1, P2, and P7 in the 2D ^1^H– ^1^H NOESY spectra. Peptide 2 displays a well-dispersed pattern of Hα resonances, reflecting distinct local environments and the presence of specific intraresidual and sequential interactions. The remaining peptides P1 and P7 show heterogenous and overlapping signals, consistent with disordered conformations in solution. **b**, Comparison of the aliphatic regions (1.0–2.6 ppm) of P1, P2, and P7 in the 2D ^1^H– ^1^H NOESY spectra. P2 exhibits a diverse set of cross-peaks corresponding to β- and γ-protons and methyl groups of side chains, suggesting a spatial arrangement exists. In contrast, the spectra of P1 and P7 show extensive overlap and reduced signal dispersion or complete lack of signals, consistent with higher conformational flexibility and lack of stable defined structure.

**Extended Data Figure 6.**
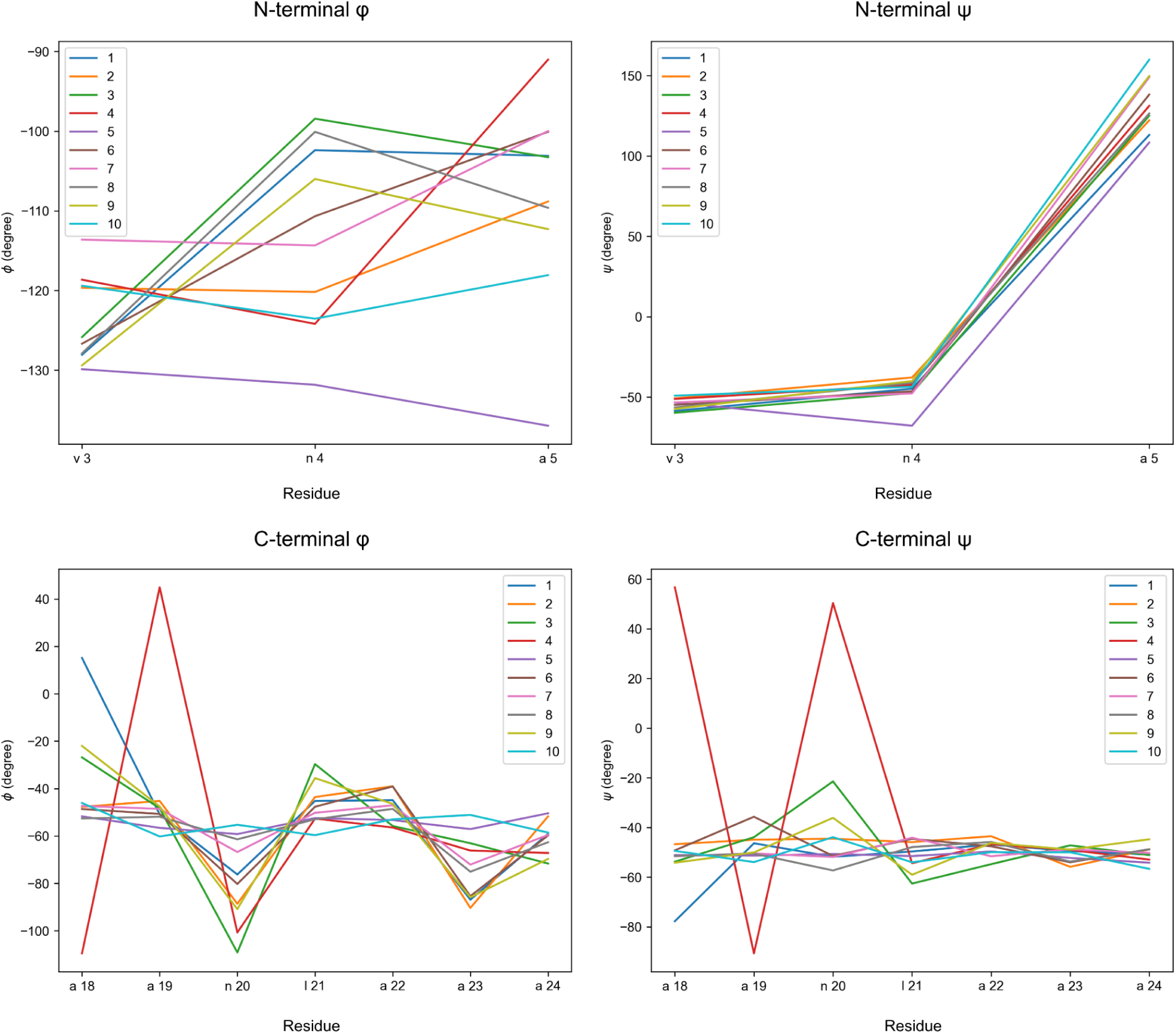
Phi and psi dihedral angles for N- and C-terminal predicted secondary structures. Analysis conducted on 10 representative structures from 10 independent replicas in MD (shown each as a different color). The three amino acids closest to the N-terminus and seven amino acids closest to the C-terminus are depicted, as described in the text.

**Extended Data Figure 7.**
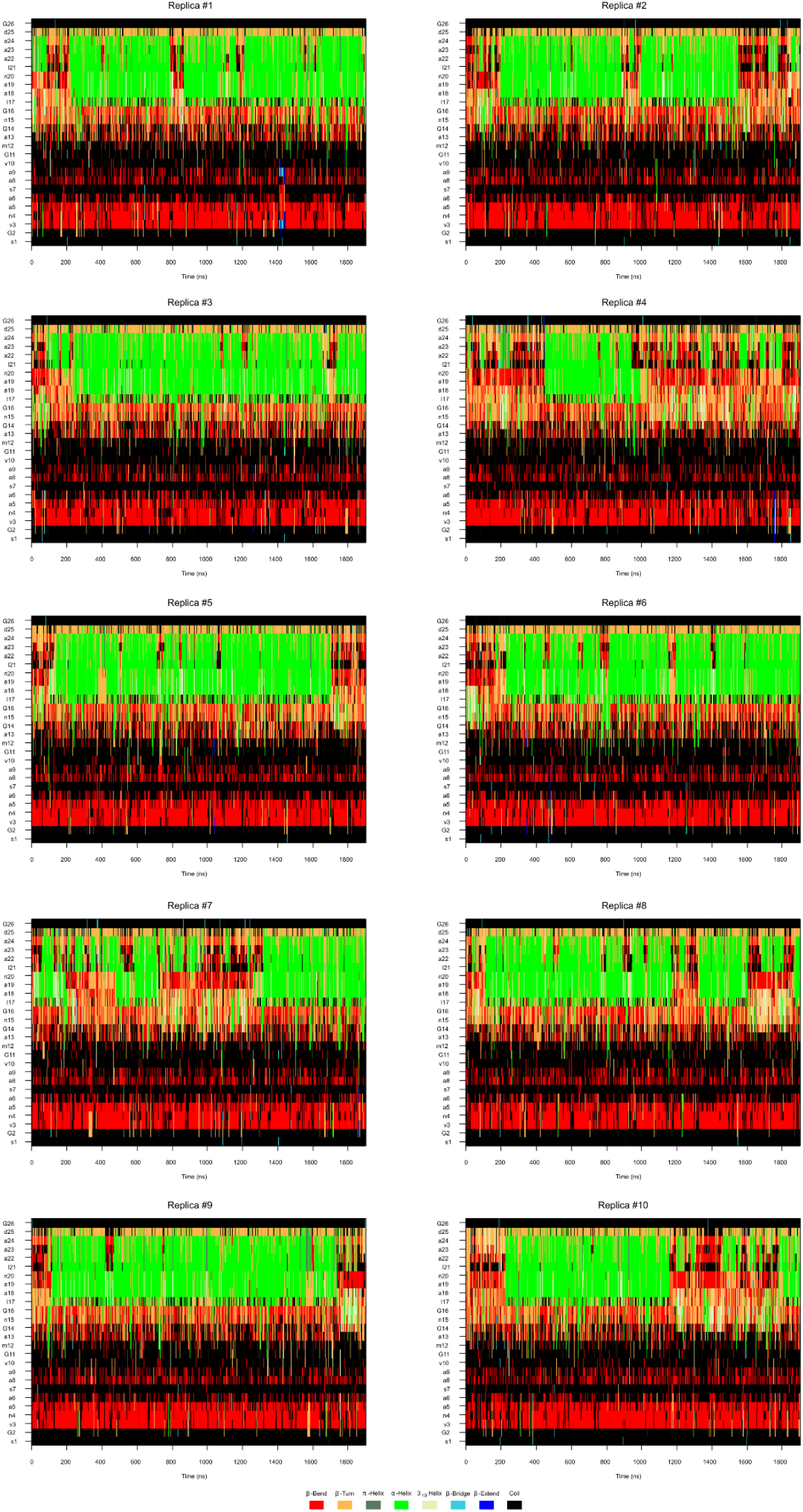
MD trajectories of P2. Per-residue evolution of P2 secondary structures in ten replicas in the course of 1.9 µs molecular dynamics. β-bend structures (red), β-turn structures (orange), coils (black), and α-helix (green) are primarily observed while π-helix (grey), 3_10_-helix (tan), β-bridge (cyan), and β-extended structures (blue) are observed less frequently.

**Extended Data Table 1.** JChem Estimated pK_A_s of Pho, Ccs, Gla, Dmg, Asp, and Glu.

| Amino Acid | JChem $pK_A$<br>Value(s) |
| --- | --- |
| Asp<sup>a</sup> | 3.82 |
| Glu<sup>a</sup> | 4.28 |
| Pho | 8.39, 1.72 |
| Ccs | 4.05 |
| Gla | 6.01, 2.31 |
| Dmg | 4.61 |

<sup>a</sup>For Asp and Glu, experimental $pK_A$ values are 3.90 and 4.07, resp.

**Extended Data Table 2.** Final 10 Alphabets LSA and x̄ EMD Values. Xeno 1 and Xeno 2 alphabets (X1, X2) underscored in bold.

| Alphabet | LSA Values | LSA Rank | EMD Values | EMD Rank |
| --- | --- | --- | --- | --- |
| **X1** | **13.8** | **4** | **3.42** | **9** |
| A2 | 20.7 | 9 | 2.06 | 1-2 |
| A3 | 17.0 | 7 | 4.28 | 10 |
| **X2** | **14.6** | **5** | **2.35** | **3** |
| A5 | 12.4 | 2 | 2.83 | 7 |
| A6 | 15.1 | 6 | 2.86 | 8 |
| A7 | 12.9 | 3 | 2.06 | 1-2 |
| A8 | 18.2 | 8 | 2.67 | 5 |
| A9 | 20.7 | 10 | 2.66 | 4 |
| A10 | 11.8 | 1 | 2.75 | 6 |

**Extended Data Table 3.**
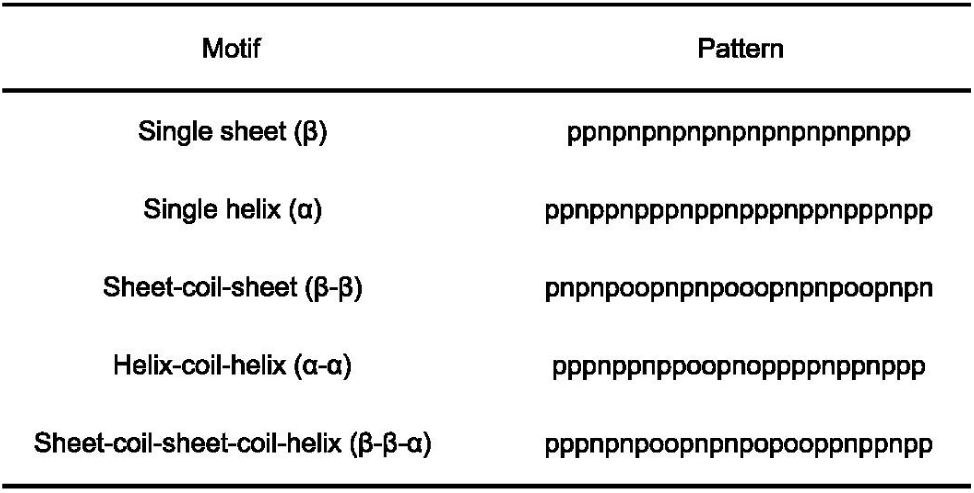
Patterns for peptide design. Polar residues are denoted in p, nonpolar are denoted by n, other residues are denoted by o.

