## Supplementary figures and images for "Xeno amino acid alphabets form peptides with familiar secondary structure"

### peptide1_spectrum.png

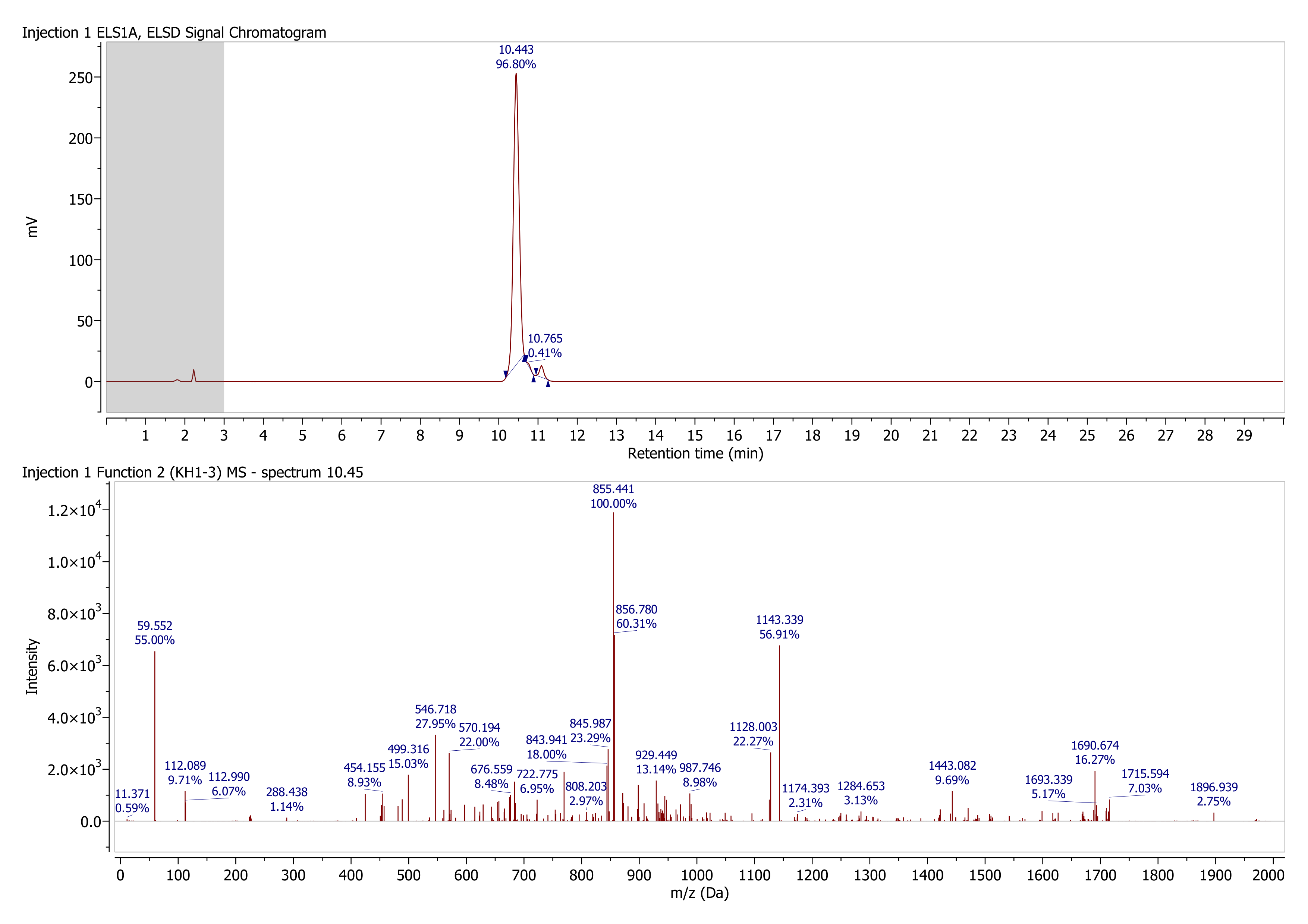

### peptide1_structure.png

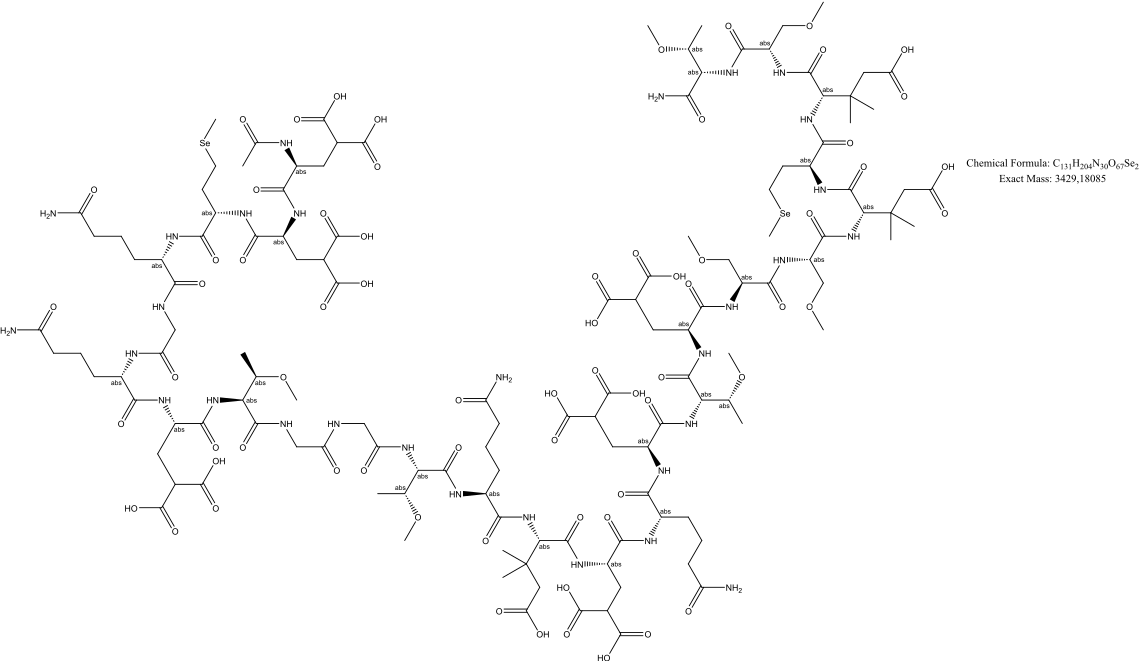

### peptide2_spectrum.png

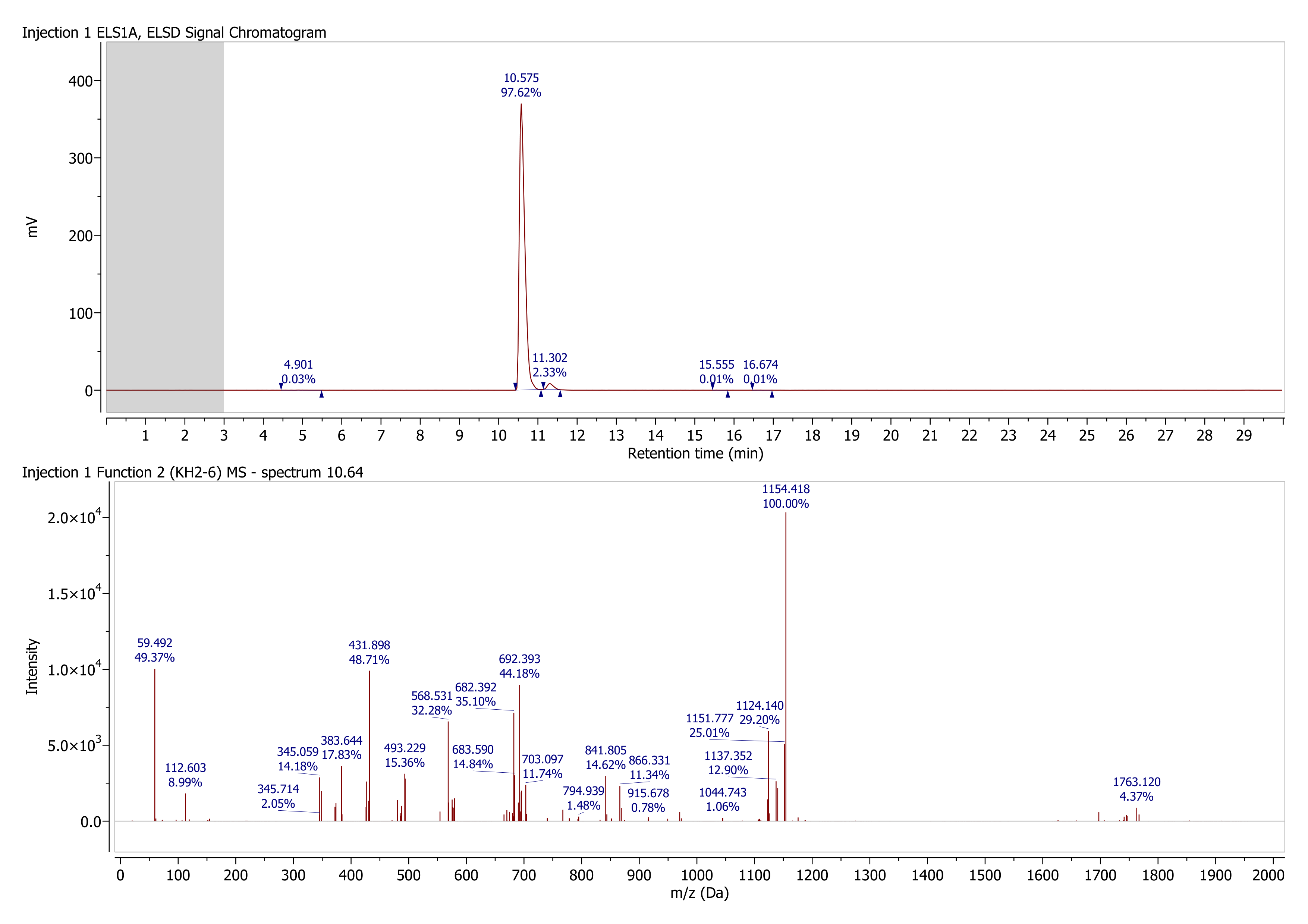

### peptide2_structure.png

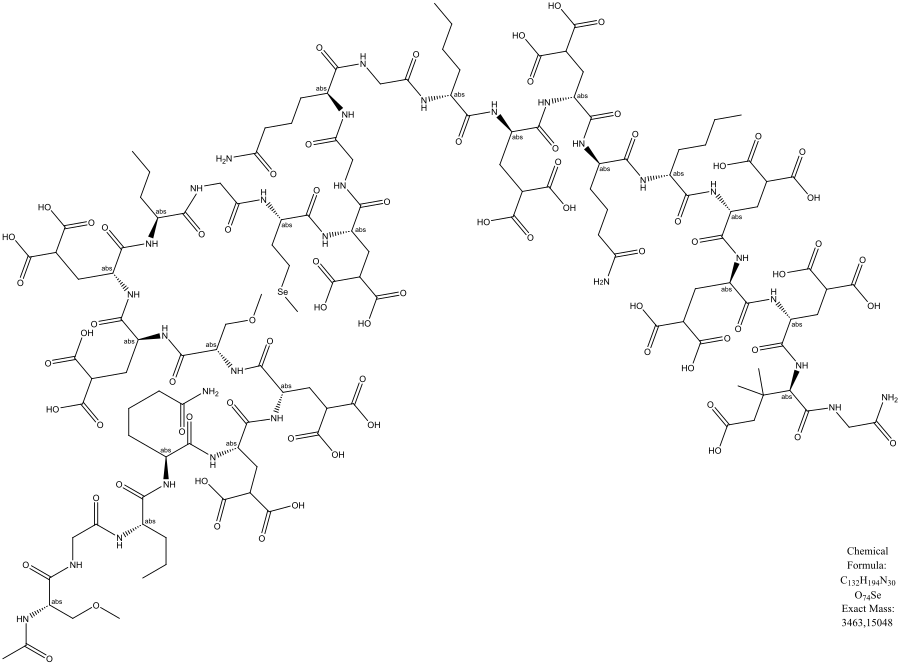

### peptide3_spectrum.png

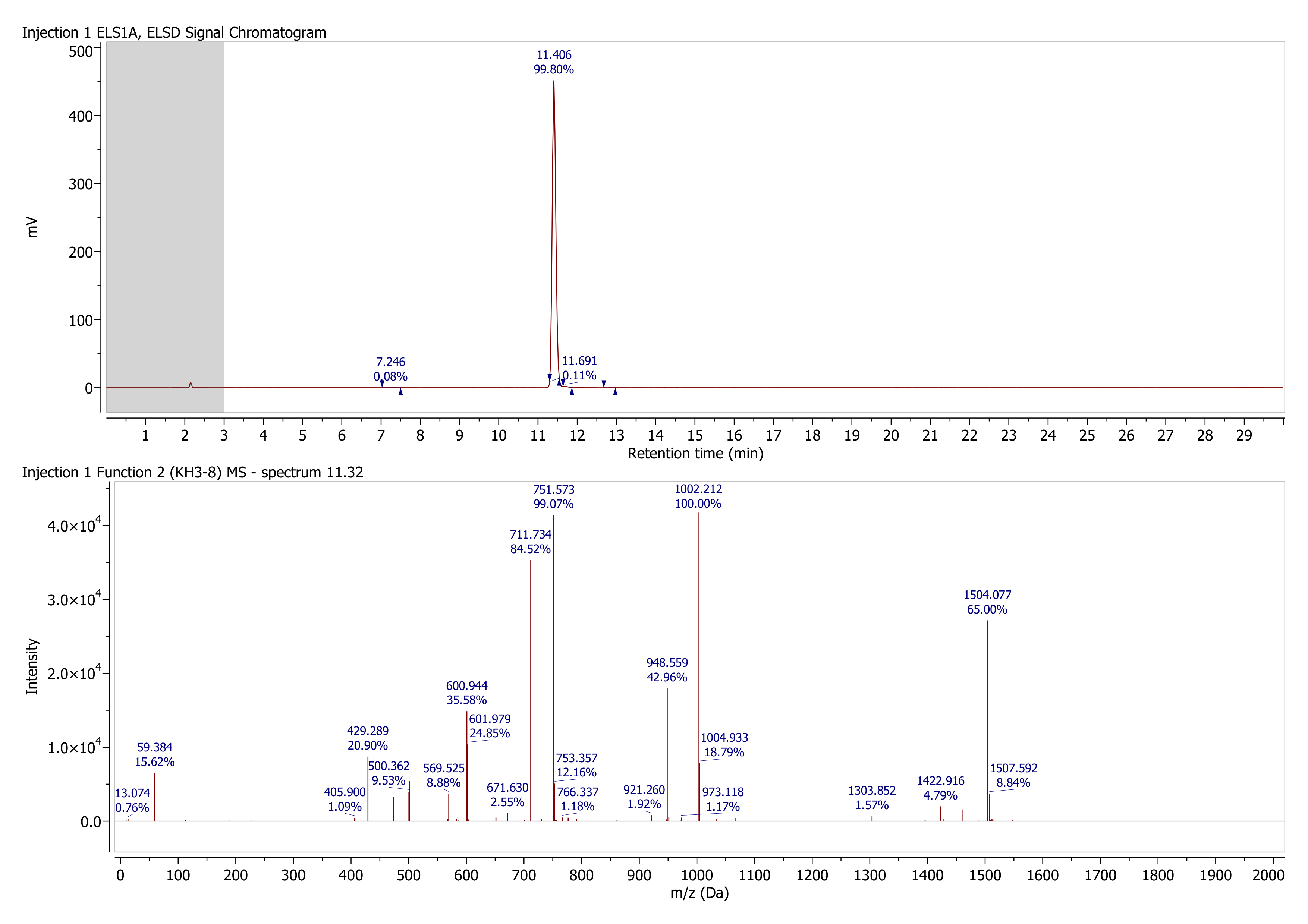

### peptide3_structure.png

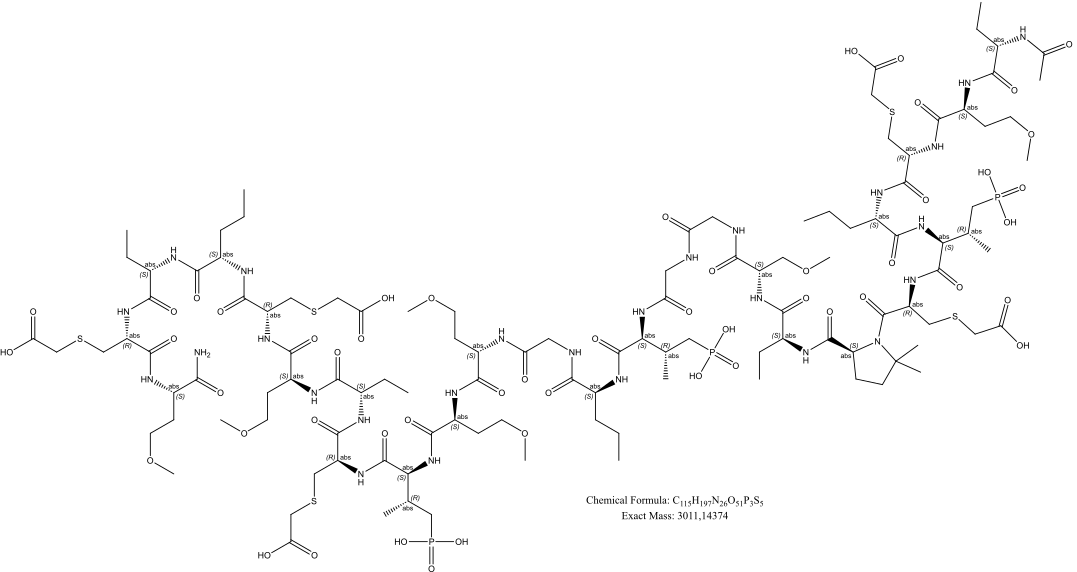

### peptide4_spectrum.png

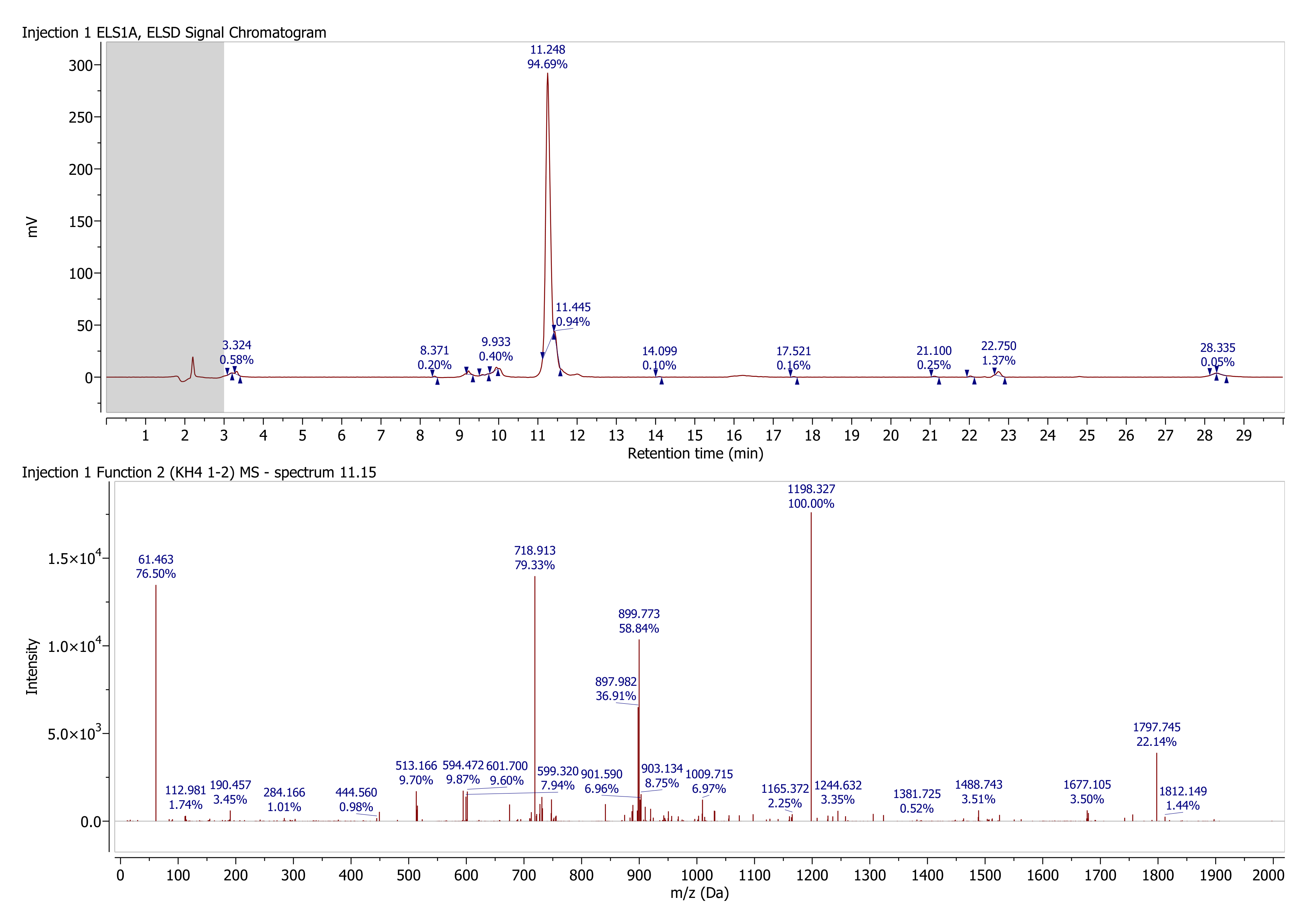

### peptide4_structure.png

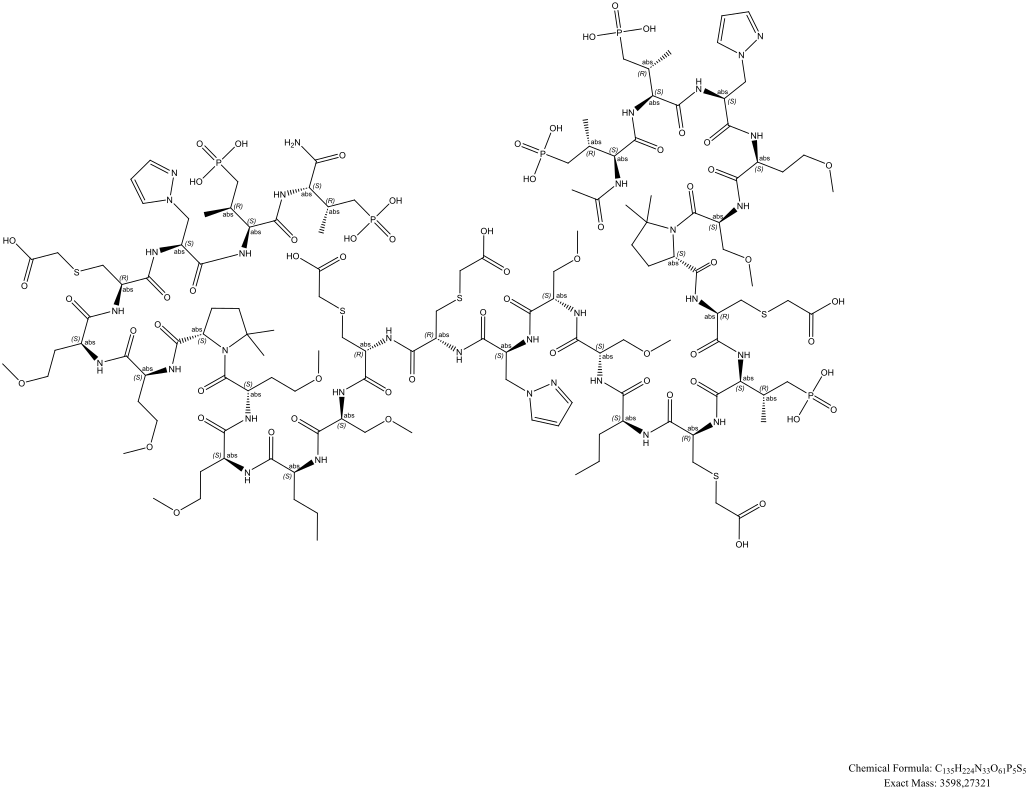

### peptide5_spectrum.png

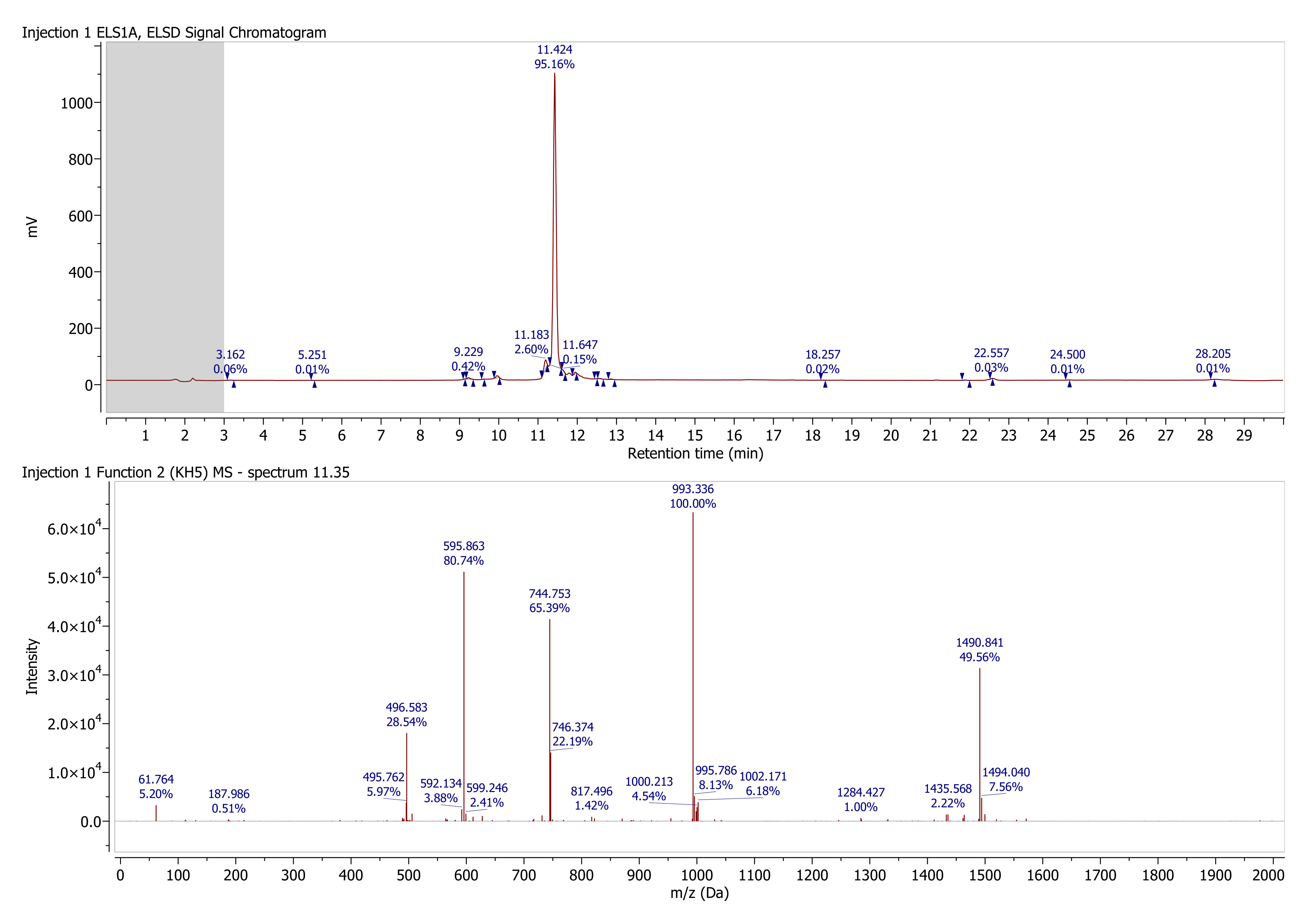

### peptide5_structure.png

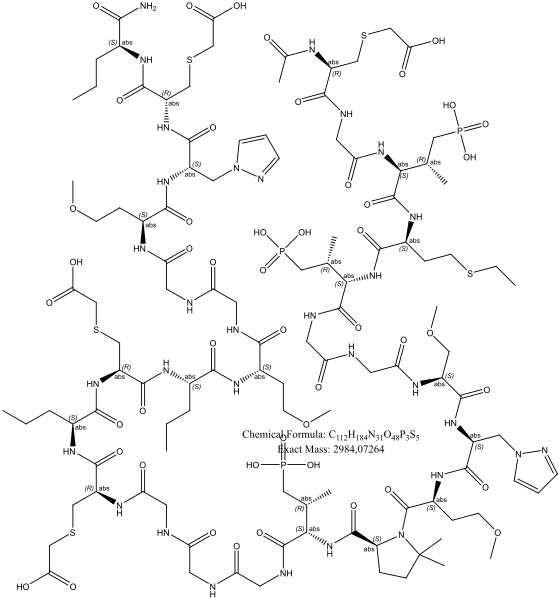

### peptide6_spectrum.png

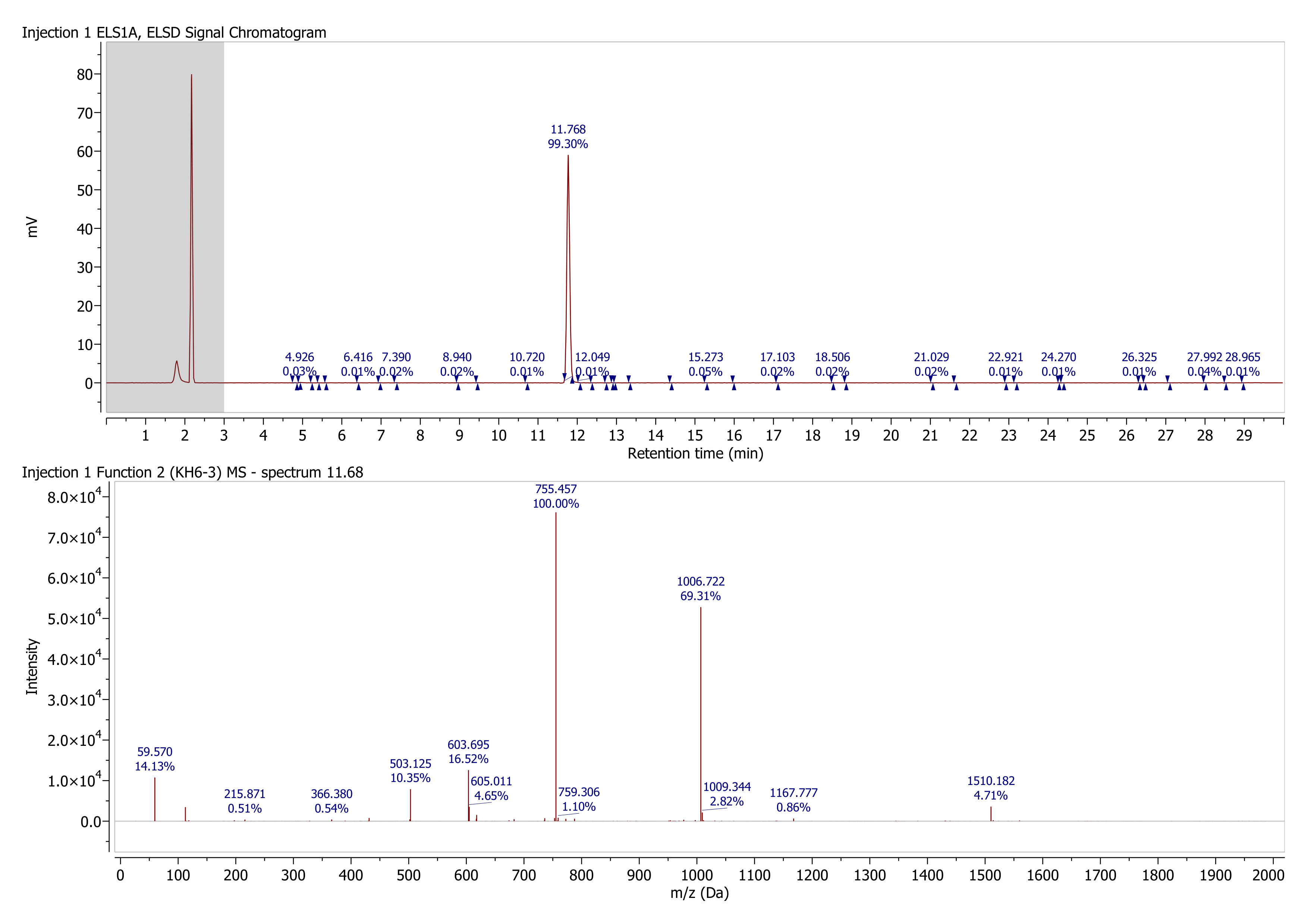

### peptide6_structure.png

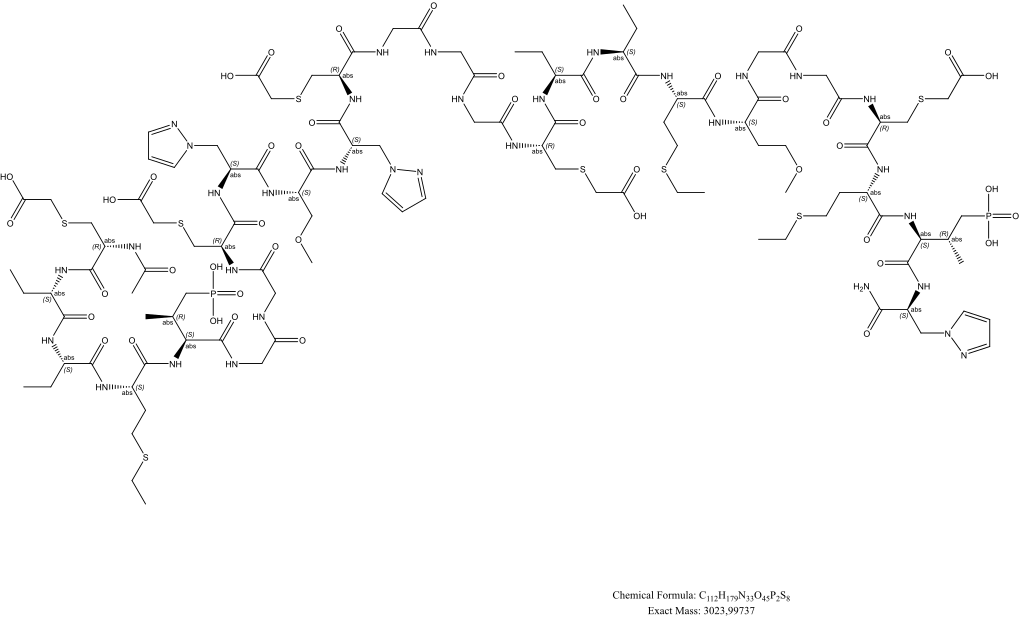

### peptide7_spectrum.png

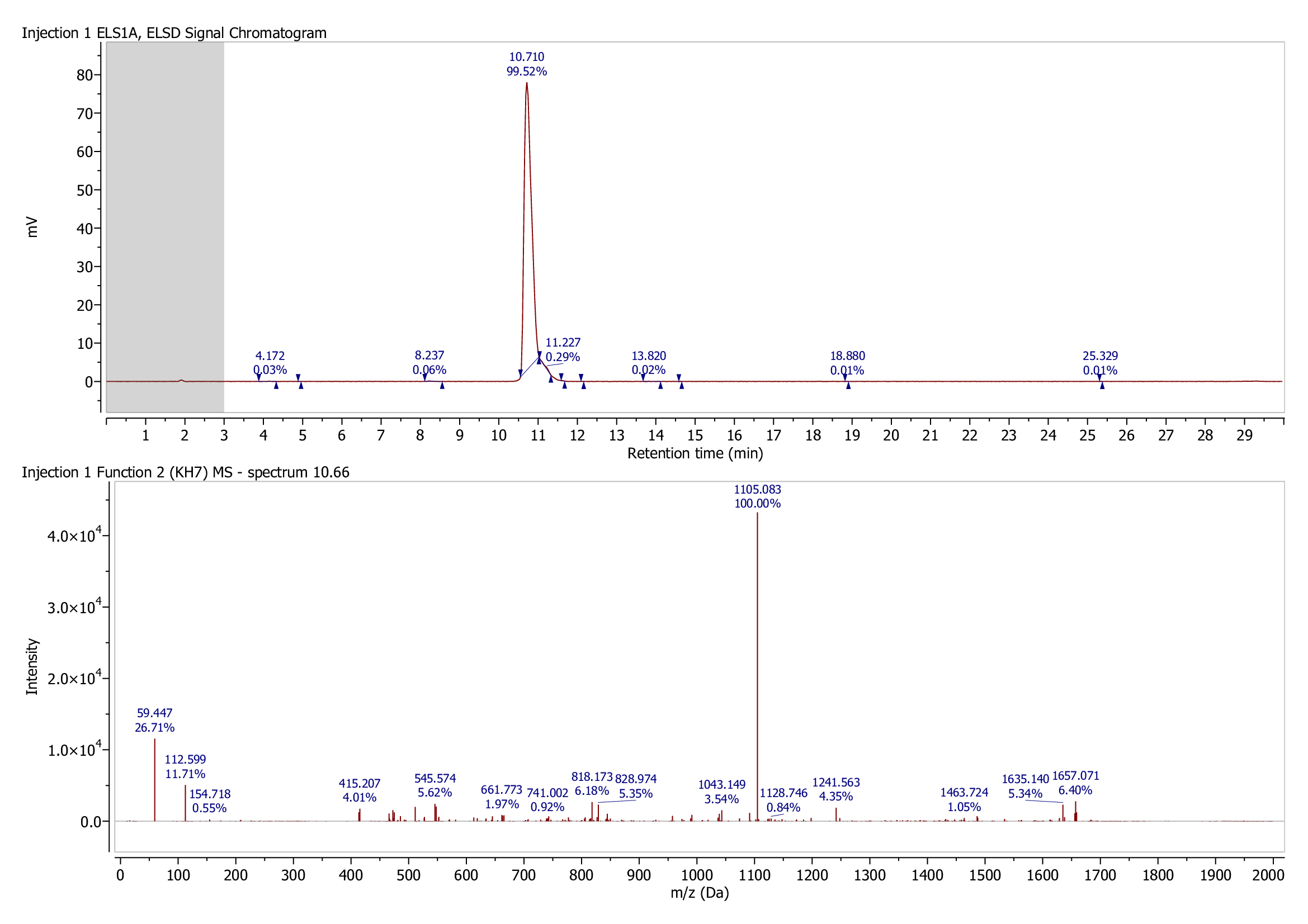

### peptide7_structure.png

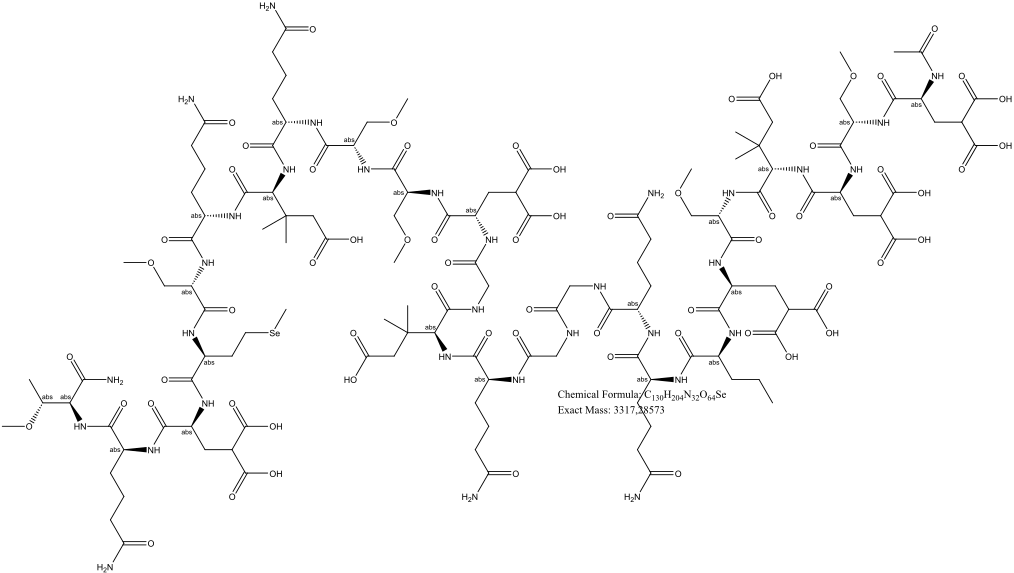
