## Supplementary material for "Xeno amino acid alphabets form peptides with familiar secondary structure": SI.

### TABLE OF CONTENTS

**Supplementary Methods** **3**
SI.1 Introduction to Coverage Theory 3
SI.2 Conformational Sampling of Tripeptides 4
SI.3 CD Spectroscopy at Various pH and TFE Addition 5
**Supplementary Data** **5**
SI.4 Synthesis Results 5
SI.5 SRCD Data 5
**Supplementary Information References** **6**

### Supplementary Methods

#### SI.1 Introduction to Coverage Theory

A longstanding approach within evolutionary biology to identify and understand adaptations is to ask: what is unusual (distinct from random) about the phenomenon under consideration?[S1–3] Previous literature[S4–7] has identified that the alphabet of 20 genetically encoded amino acids exhibits a statistically unusual profile in terms of van der Waals' volume (size) and LogP (hydrophobicity). This profile, termed "*Coverage*" has been defined as the range of values exhibited by a set of amino acids (calculated as the sum of the intervals between rank-ordered amino acids), and the evenness with which members of the alphabet distribute across this range[S4] (calculated as the sample variance of these intervals, where a larger evenness value indicates a less even distribution - see Extended Data Fig. 1a).

Hydrophobicity and volume profoundly affect protein folding. Hydrophobic collapse is accepted as an underlying principle[S8–10] and perhaps even the driver[S11] of protein folding. Volume determines steric constraints limiting structure formation, such as $\varphi/\psi$ torsion angles[S12] and core-packing within proteins[S13]. The statistically unusual properties of life's amino acid alphabet in these two physicochemical properties therefore links plausibly to their evolutionary success as a set[S14].

Previously, this pattern has been investigated[S15] for the E10 amino acid alphabet (G, A, D, V, S, E, P, L, T, I). Statistical nonrandomness is evident though not as strong for the early ten as it is for C20. Our study therefore uses heuristic search to design two xeno amino acid alphabets (Xeno 1 and Xeno 2) which emulate the physicochemical coverage of E10.

#### SI.2 Conformational Sampling of Tripeptides

For the purpose of sequence design augmentation, we wanted to explore the conformational space of tripeptides which can be constructed using amino acids of both Xeno alphabets. As the systematic sampling of PES used for single amino acids in the main text would be computationally infeasible for tripeptides due to the high demand for computational time (as the number of conformers for a single tripeptide would start at 2,985,984 in the approach used for amino acids), we run CREST to obtain the local minima on PES only. CREST was run in the same manner as for amino acids (see methods in the main text), again with the BP86-DB(BJ)/DGauss-DZVP//COSMO-RS single point, but with no constraints on backbone in order to freely explore PES. The RMSD threshold (*--rthr*) was set to 1. All other settings were default. Final conformational energies were again calculated like for the amino acids.

For all conformers of each tripeptide, the secondary structure (alpha-helix, extended, polyproline II, other) of each amino acid was determined and redundant conformers were removed in the same manner as in the previous work[S16]. We then calculated the dG(helix-extended) window as the energy difference between lowest all-alpha-helical and all-extended conformer. Finally, top 20 tripeptides with dG(helix-extended) > 3 kcal·mol$^{-1}$ and < -3 kcal·mol$^{-1}$, excluding tripeptides containing glycine, were designated as pro-alpha-helical and pro-extended tripeptides, respectively, and used for augmentation (see “SI_Sequences.xlsx” for a complete list).

#### SI.3 CD Spectroscopy at Various pH and TFE Addition

Secondary structure analysis was performed by far-UV CD spectroscopy. The far-UV CD spectra were recorded using a Chirascan-plus spectrophotometer (Applied Photophysics, Leatherhead, UK) over the wavelength range 190–260 nm in steps of 1 nm with an averaging time of 1 s per step. Cleared samples at 0.2 mg/ml nominal concentration in 1 mm path-length quartz cells were placed into a cell holder, and spectra were recorded at room temperature. The CD signal was obtained as ellipticity in units of millidegrees, and the resulting spectra were averaged from two scans and buffer-spectrum was subtracted. All CD measurements were repeated twice as replicates.

The effect of pH on secondary structure was estimated by diluting stock solutions of peptide libraries with 10 mM ABP buffers at pH 3.0, 5.0, 7.4, 9.0, and 11.0. The effect of 2,2,2-trifluoroethanol on secondary structure was estimated by diluting stock solutions of peptide libraries with a series of 10 mM ABP buffer (pH 7.4) containing 0–90% (v/v) 2,2,2-trifluoroethanol. The peptide solutions were gently mixed at room temperature for 30 min and then centrifuged at 21,300 *g* for 15 min at 4 °C in order to remove the insoluble part.

### Supplementary Data

#### SI.4 Synthesis Results

The synthesis data of peptide 1-7 are deposited in the "SI_Peptide_1-7_Synthesis" folder as part of the Supplementary Data.

#### SI.5 SRCD Data

The synchrotron radiation circular dichroism spectra of all measured peptides are deposited in .xlsx format in "SI_SRCD.xlsx" as part of the Supplementary Data.

### Supplementary Information References


S1. Darwin, C. *On the Various Contrivances by Which British and Foreign Orchids Are Fertilised by Insects: And on the Good Effect of Intercrossing*. (Cambridge University Press, Cambridge, 2011). doi:10.1017/CBO9780511910197.

S2. Parker, G. A. & Smith, J. M. Optimality theory in evolutionary biology. *Nature* **348**, 27–33 (1990).

S3. Freeland, S. J. & Hurst, L. D. The Genetic Code Is One in a Million. *J. Mol. Evol.* **47**, 238–248 (1998).

S4. Philip, G. K. & Freeland, S. J. Did Evolution Select a Nonrandom “Alphabet” of Amino Acids? *Astrobiology* **11**, 235–240 (2011).

S5. Ilardo, M., Meringer, M., Freeland, S., Rasulev, B. & Cleaves II, H. J. Extraordinarily Adaptive Properties of the Genetically Encoded Amino Acids. *Sci. Rep.* **5**, 9414 (2015).

S6. Mayer-Bacon, C., Meringer, M., Havel, R., Aponte, J. C. & Freeland, S. A Closer Look at Non-random Patterns Within Chemistry Space for a Smaller, Earlier Amino Acid Alphabet. *J. Mol. Evol.* **90**, 307–323 (2022).

S7. Brown, S. M., Voráček, V. & Freeland, S. What Would an Alien Amino Acid Alphabet Look Like and Why? *Astrobiology* **23**, 536–549 (2023).

S8. Agashe, V. R., Shastry, M. C. R. & Udgaonkar, J. B. Initial hydrophobic collapse in the folding of barstar. *Nature* **377**, 754–757 (1995).

S9. Kauzmann, W. Some Factors in the Interpretation of Protein Denaturation1. in *Advances in Protein Chemistry* (eds Anfinsen, C. B., Anson, M. L., Bailey, K. & Edsall, J. T.) vol. 14 1–63 (Academic Press, 1959).

S10. Robson, B. & Pain, R. H. Analysis of the code relating sequence to conformation in proteins: Possible implications for the mechanism of formation of helical regions. *J. Mol. Biol.* **58**, 237–257 (1971).

S11. Salamatova, E. *et al.* Hydrophobic Collapse in N-Methylacetamide–Water Mixtures. *J. Phys. Chem. A* **122**, 2468–2478 (2018).

S12. DasGupta, D., Kaushik, R. & Jayaram, B. From Ramachandran Maps to Tertiary Structures of Proteins. *J. Phys. Chem. B* **119**, 11136–11145 (2015).

S13. Lim, W. A. & Sauer, R. T. The role of internal packing interactions in determining the structure and stability of a protein. *J. Mol. Biol.* **219**, 359–376 (1991).

S14. Mayer-Bacon, C., Agboha, N., Muscalli, M. & Freeland, S. Evolution as a Guide to Designing xeno Amino Acid Alphabets. *Int. J. Mol. Sci.* **22**, 2787 (2021).

S15. Mayer-Bacon, C. & Freeland, S. J. A broader context for understanding amino acid alphabet optimality. *J. Theor. Biol.* **520**, 110661 (2021).

S16. Osifová, Z. *et al.* What are the minimal folding seeds in proteins? Experimental and theoretical assessment of secondary structure propensities of small peptide fragments. *Chem. Sci.* **15**, 594–608 (2024).
